# BPTF cooperates with MYCN and MYC to link neuroblastoma cell cycle control to epigenetic cellular states

**DOI:** 10.1101/2024.02.11.579816

**Authors:** Irene Felipe, Jaime Martínez-de-Villarreal, Khushbu Patel, Jorge Martínez-Torrecuadrada, Liron D. Grossmann, Giovanna Roncador, Mariona Cubells, Alvin Farrell, Nathan Kendsersky, Sergio Sabroso-Lasa, Lucía Sancho-Temiño, Kyabeth Torres, Daniel Martinez, Javier Muñoz Perez, Fernando García, Jenny Pogoriler, Lucas Moreno, John M. Maris, Francisco X. Real

**Author notes:** Correspondence should be addressed to: Francisco X. Real, Centro Nacional de Investigaciones Oncológicas-CNIO Melchor Fernández Almagro, 3, 28029-Madrid, Spain.

## Abstract

The nucleosome remodeling factor BPTF is required for the deployment of the MYC-driven transcriptional program. Deletion of one *Bptf* allele delays tumor progression in mouse models of pancreatic cancer and lymphoma. In neuroblastoma, MYCN cooperates with the transcriptional core regulatory circuitry (CRC). High *BPTF* levels are associated with high-risk features and decreased survival. BPTF depletion results in a dramatic decrease of cell proliferation. Bulk RNA-seq, single-cell sequencing, and tissue microarrays reveal a positive correlation of *BPTF* and CRC transcription factor expression. Immunoprecipitation/mass spectrometry shows that BPTF interacts with MYCN and the CRC proteins. Genome-wide distribution analysis of BPTF and CRC in neuroblastoma reveals a dual role for BPTF: 1) it co-localizes with MYCN/MYC at the promoter of genes involved in cell cycle and 2) it co-localizes with the CRC at super-enhancers to regulate cell identity. The critical role of BPTF across neuroblastoma subtypes supports its relevance as a therapeutic target.

## Introduction

Cancer genome projects have unveiled a crucial role of genes coding for proteins involved in chromatin activity in tumor development ^1, 2^. Many of these proteins have enzymatic activity towards histone and non-histone proteins. Others do not have any known enzymatic activity, yet there is increasing evidence that they play a role in tumorigenesis. Among them is BPTF, a 3046-residue component of the NURF complex that hydrolyses ATP to induce nucleosome sliding within chromatin. BPTF contains 2 PHD finger domains involved in methylated lysine recognition and a Bromodomain (BrD) that binds acetylated lysines, epigenetic marks present in histones. BPTF provides specificity to NURF chromatin binding through its interaction with histone modifications in transcriptionally active genes (i.e., H3K4me3, H4K16Ac, and H2A.Z) ^3, 4, 5, 6^. BPTF cooperates with CTCF to regulate gene expression ^7^ and is required for early embryonic development ^8^.

Our group identified *BPTF* as a novel gene involved in several tumor types including pancreatic cancer, urothelial bladder cancer, and lymphoma ^9, 10, 11^. We have reported that BPTF interacts directly with MYC (c-MYC), as demonstrated by co-immunoprecipitation (co-IP) and proximity ligation assays (PLA) ^10^. This interaction is critical for MYC function: *BPTF* knockdown leads to a decrease in MYC binding to DNA and impaired activation of the MYC transcriptional programme. These molecular effects are accompanied by a robust inhibition of cellular proliferation *in vitro*. We have assessed the effects of *Bptf* inactivation *in vivo* using genetic mouse models of pancreatic cancer and lymphoma ^11^. In both systems, deletion of one *Bptf* allele is sufficient to delay tumor initiation and increase mouse survival. These results highlight that a reduction of BPTF function is sufficient to induce antitumor effects. In addition, it has been proposed that BPTF inhibition leads to increased sensitivity to antitumor therapies ^12,13^ and work to establish small molecule inhibitors is ongoing ^14^. However, little is known about the molecular mechanisms of action of BPTF to promote tumorigenesis.

Neuroblastoma is an extra-cranial solid tumor originating from undifferentiated neural crest-derived cells that is widely metastatic in 50% of patients at diagnosis and continues to show one of the least favorable outcomes among pediatric cancers ^15, 16, 17^. *MYCN* amplification is strongly associated with poor outcome and is integrated into risk stratification and treatment planning ^18^. Neuroblastomas exhibit epigenetically mediated cellular plasticity with a highly proliferative adrenergic (ADRN) cell state that can transdifferentiate into a more quiescent and therapy resistant mesenchymal (MES) cell state ^19^. These cell states are maintained by a group of ADRN- and MES-specific transcription factors that form the so-called Core Regulatory Circuit (CRC). MYCN regulates the activity of the CRC transcription factors, including GATA3, PHOX2B, HAND2, ISL1, and TBX2, to maintain ADRN cell identity at the level of distal super enhancers ^20, 21, 22, 23, 24^. Loss of lineage identity and acquisition of a MES program have been shown to characterize neuroblastoma progression and therapeutic resistance, highlighting this aspect of tumor biology. Several transcription factors, including PRRX1, NOTCH, TWIST, FOSL1, FOSL2, and RUNX1, have been identified as drivers of the MES phenotype through enhancer reprogramming ^22, 25, 26^.

We hypothesized that BPTF plays a key cooperative role in high-risk neuroblastoma tumorigenesis based on the following: i) *MYCN* amplification is a major oncogenic driver in this tumor; ii) tumors lacking *MYCN* amplification often display high-level MYC expression; iii) BPTF might also be required for MYCN transcriptional activity; and iv) the *BPTF* locus lies on chr. 17q24, a region that shows copy number gain in the vast majority of high-risk neuroblastomas, as well as in other tumors ^27, 28, 29^.

Here, we follow on the observation that *BPTF* expression in human neuroblastoma is highest among all adult and pediatric cancers and focus on the association of *BPTF* expression with neuroblastoma clinical features and patient outcomes. We provide evidence linking BPTF to the activity of both MYC proteins and the neuroblastoma CRC and perform an in-depth analysis of the comparative genome-wide distribution of BPTF, MYC proteins, and CRC components in neuroblastoma cells. While BPTF binds preferentially to gene promoters, it is also present in super-enhancer regions where it co-binds with the CRC members and with MYCN/MYC. Finally, we show that BPTF expression is dynamically regulated during the neuroblastoma ADRN-MES transition. Altogether, our work demonstrates that BPTF plays a key role in neuroblastoma, being involved in the regulation of cell cycle and in epigenetically-regulated cell identity programs by modulating promoter and super-enhancer activity, respectively, and it represents a possible therapeutic target.

## RESULTS

### High *BPTF* expression levels are associated with high-risk and overall survival

Analysis of the RNA-seq data from the UCSC’s Treehouse Childhood Cancer Initiative (https://treehousegenomics.ucsc.edu), which includes childhood tumors, showed that neuroblastoma ranks third regarding highest *BPTF* mRNA levels among all human cancers (**Figure 1A**), suggesting its biological relevance in this tumor. Similar analysis of the Cancer Cell Line Encyclopedia (CCLE) revealed that neuroblastoma cell lines express the highest levels of *BPTF* among all tumor types analyzed (**Figure 1B**) and we have confirmed high expression using RT-qPCR (**Figure 1C**). These results indicate that neuroblastoma cell lines are suitable models to assess its biological role. BPTF detection at the protein level is limited by the lack of high-quality antibodies that can be used in a diverse set of applications. To overcome this problem, we developed and validated new mouse monoclonal antibodies recognizing BPTF using western blotting (WB) and immunohistochemistry (IHC) on formalin-fixed paraffin-embedded (FFPE) neuroblastoma cell pellets (**Figures 1D and S1**). Specificity of the antibodies was confirmed in both types of assays using cells in which BPTF was knocked down through lentiviral shRNA transduction. Lentiviral knockdown of *BPTF* markedly reduced the proliferation of a wide range of neuroblastoma cells: KELLY, a prototypic ADRN line, was exquisitely sensitive and extensive cell death occurred over 24-48h; similar effects were observed on SK-N-AS, a MES line, over 72-96h (**Figure 1D, E**).

**Figure 1.**
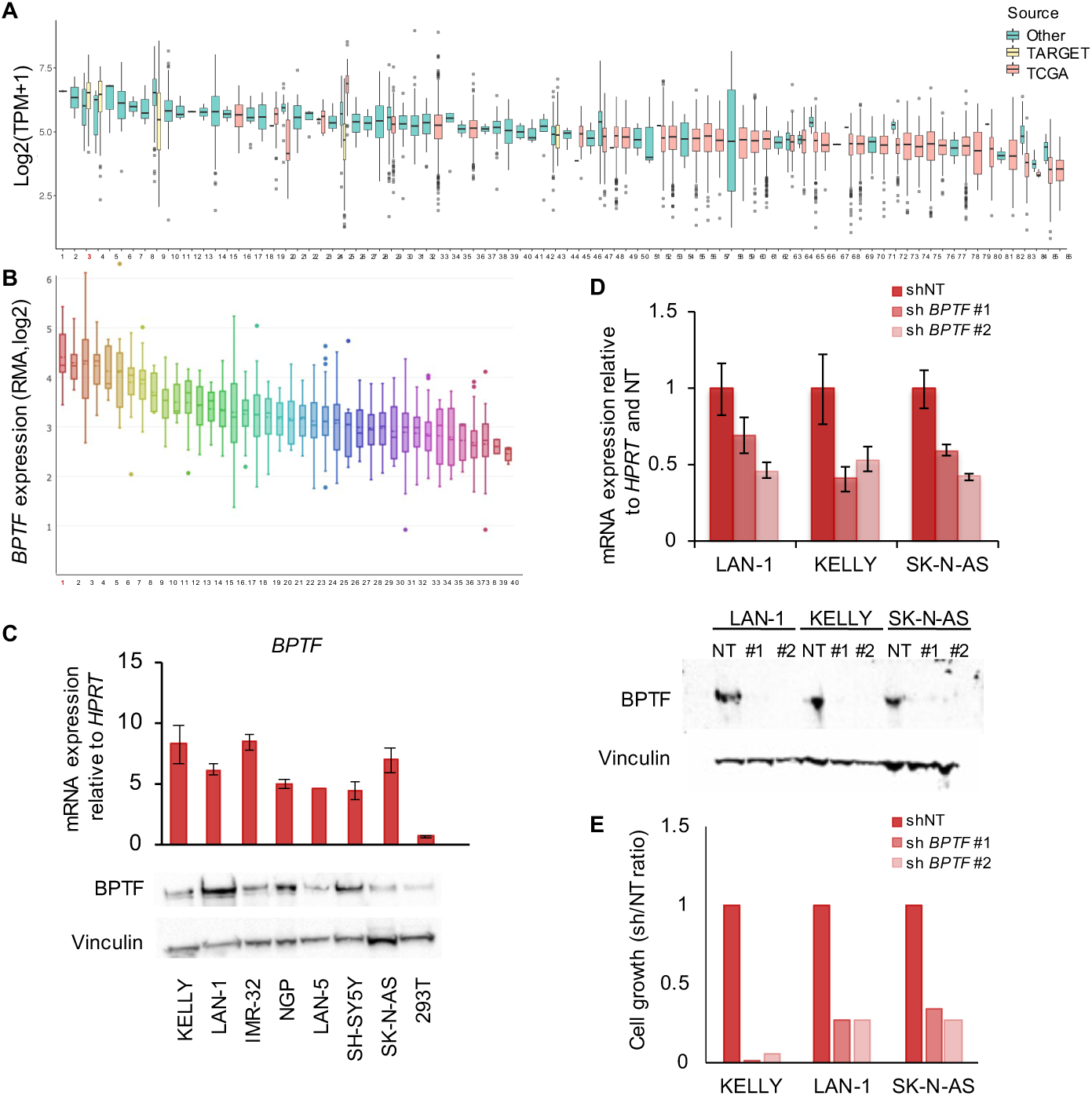
*BPTF* is expressed at high levels in neuroblastoma and its knockdown has antiproliferative effects, regardless of *MYCN* status. (A) *BPTF* expression across cancers: data from the UCSC Treehouse Dataset (https://treehousegenomics.ucsc.edu), ordered by decreasing *BPTF* levels. (1, Rb; 2, ET; **3, Neuroblastoma;** 4, Wilms tumor; 5, Meningioma; 6, CPC; 7, Neurofibroma; 8, MEC; 9, ALL; 10, EPN; 11, DSRCT; 12, ULS; 13, IFS; 14, AMKL; 15, GCT; 16, ESCA; 17, DNET; 18, Leukemia; 19, TGTC; 20, SS; 21, Supratentorial embryonal tumor NOS; 22, IMT; 23, MPNST; 24, GIST; 25, LAML; 26, UPS NOS; 27, Undifferentiated spindle cell SARC; 28, SCLC; 29, Glioma; 30, ES; 31, EMRS; 32, Lymphoma; 33, BRCA; 34, Spindle cell/sclerosing rhabdomyosarcoma; 35, RT; 36, STAD; 37, ALAL; 38, ARMS; 39, RMS; 40, Fibromatosis; 41, ES; 42, Acute leukemia; 43, OS; 44, JMML; 45, CHOL; 46, TC; 47, GBM; 48, LUAD; 49, UCS; 50, Melanoma; 51, NPC; 52, OV; 53, SKCM; 54, ATRT; 55, LSCC; 56, THYM; 57, MESO; 58, PNET; 59, DDLPS; 60, BLCA; 61, PRAD; 62, IMF; 63, SARC; 64, Phe & Para; 65, THCA; 66, MFS; 67, INI-deficient STS NOS; 68, KIRC; 69, CESC; 70, HBL; 71, PAAD; 72, COAD; 73, KIRP; 74, LEIO; 75, UCEC; 76, READ; 77, CML and ALL; 78, HNSCC; 79, DLBCL; 80, UPS; 81, MPN; 82, UVM; 83, ACC; 84, FHCC; 85, HCC; 86, KICH). (B) Among cell lines, neuroblastoma is one of the tumor types expressing highest levels of *BPTF* mRNA (Cancer Cell Line Encyclopedia, RNA-seq data). (**1, Neuroblastoma;** 2, Burkitt L; 3, DLBCL; 4, T-cell ALL; 5, Leukemia; 6, B-cell L; 7, B-cell ALL; 8, SCLC; 9, PNET; 10, T-cell L; 11, CML; 12, EWS; 13, MM; 14, STS; 15, AML; 16, BRCA; 17, NA; 18, STAD; 19, Endometrium; 20, PRAD; 21, Hodgkin L; 22, OS; 23, Colorectal; 24, LNSC; 25, LIHC; 26, Bile duct; 27, PAAD; 28, ESCA; 29, Glioma; 30, THCA; 31, OV; 32, Urinary tract; 33, Melanoma; 34, MESO; 35, Other; 36, ChS; 37, Upper aerodigestive; 38, Kidney; 39, Meningioma; 40, Giant cell tumor). (C) Neuroblastoma cells express high levels of *BPTF* mRNA and protein. HEK293T cells were included as negative control. (D) Lentiviral *BPTF* knockdown results in 40-60% lower levels of expression at the mRNA and protein levels, as detected with TOR249C Ab. (E) *BPTF* knockdown has anti-proliferative effects on neuroblastoma cells (data shown correspond to the 48h timepoint).

**Figure S1.**
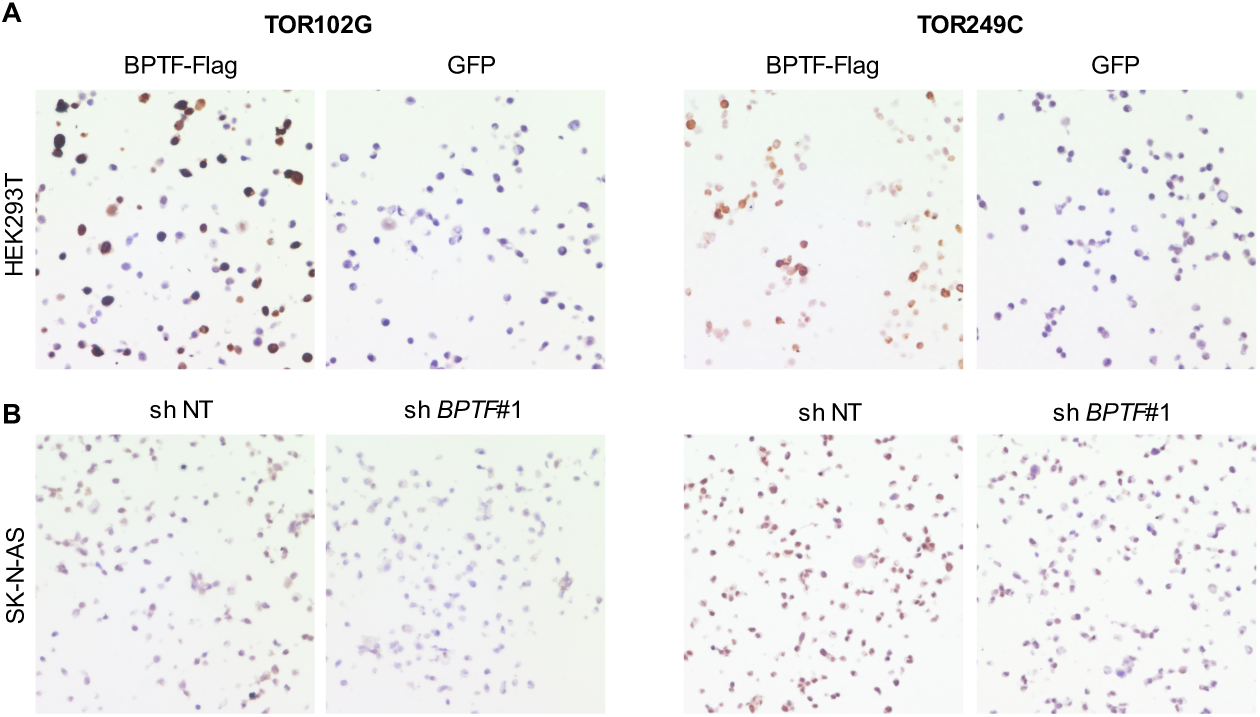
Validation of the specificity of novel anti-BPTF monoclonal antibodies. Immunohistochemical analysis of BPTF expression in cultured cells, detected with antibodies TOR102G (left) and TOR249C (right). (A) HEK293T cells overexpressing BPTF-Flag or GFP as negative control. (B) SK-N-AS neuroblastoma cells upon *BPTF* knockdown (shNT as positive control).

To assess the association between *BPTF* expression and clinical and molecular features in clinically-relevant neuroblastoma groups, we analyzed RNA-seq data from several patient cohorts. The comparison of the clinical and molecular features of the patient series analyzed is shown in **Supplementary Table 1**. First, we used the SEQC cohort ^30^, including 498 primary tumor samples, 18% of which harbor *MYCN* amplification and 35% of which are classified as high-risk. To validate the findings, we used the GMKF cohort (dbGaP phs001436.v1. p1), composed of 209 primary tumors, 26% of which are high-risk. *BPTF* was highly expressed in primary samples of both cohorts, levels being higher in males (P=0.006) and in high-risk tumors (SEQC, *P*= 6.3e-10; GMKF, *P*=0.015) (**Figure 2A and S2A**). Among the latter, *BPTF* mRNA levels were similar in cases with or without *MYCN* amplification (**Figure 2A and S2A**). High *BPTF* expression was associated with worse overall survival (OS) in the SEQC cohort (*P*=0.003) (**Figure 2B**). In multivariable analyses, *BPTF* expression levels did not emerge as significantly associated with the outcomes analyzed, likely due to the strong effect of *MYCN* amplification and risk stratification (**Figure 2C and S2B**).

**Supplementary Table 1.**
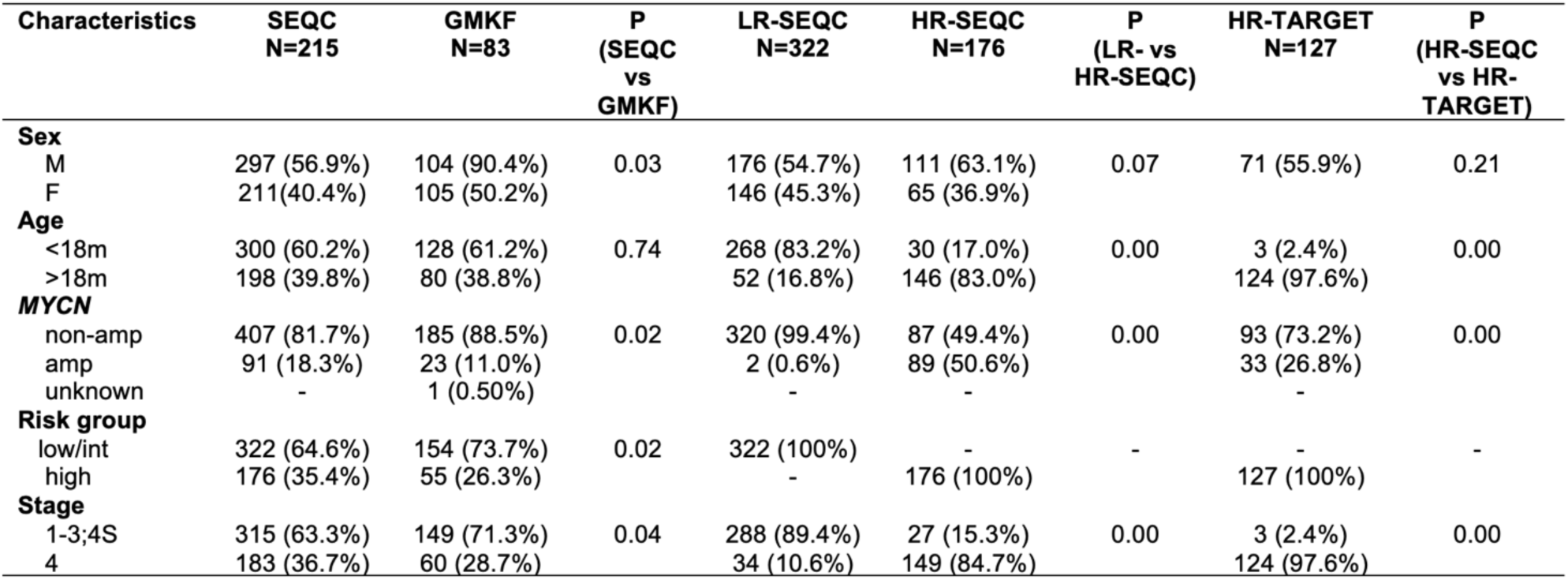
Comparison of clinical information among human neuroblastoma patient datasets.

**Figure S2.**
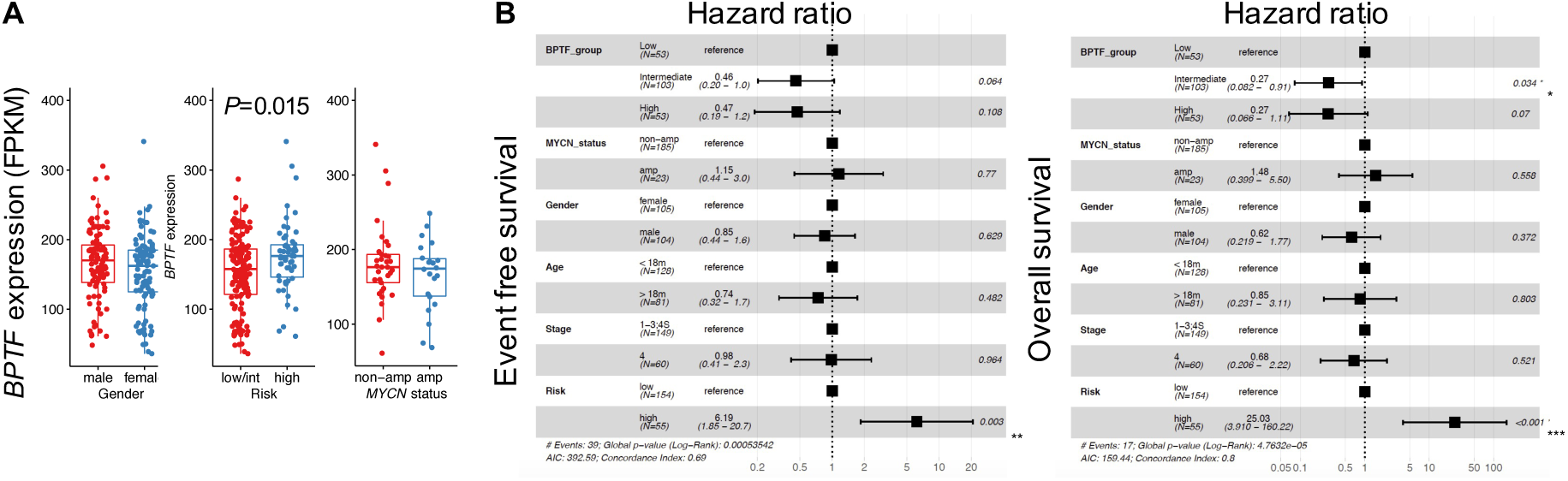
Association of *BPTF* expression with gender, risk, and *MYCN* status in the human GMKF dataset. (A) Association of *BPTF* levels and gender, risk group, and *MYCN* amplification (Wilcoxon test). (B) Cox proportional hazards regression models showing the association of relevant clinical covariates and *BPTF* expression with EFS and OS. Error bars represent the 95% confidence intervals and the indicated p values are from log-rank test. Sample size for groups is indicated on the figures.

**Figure 2.**
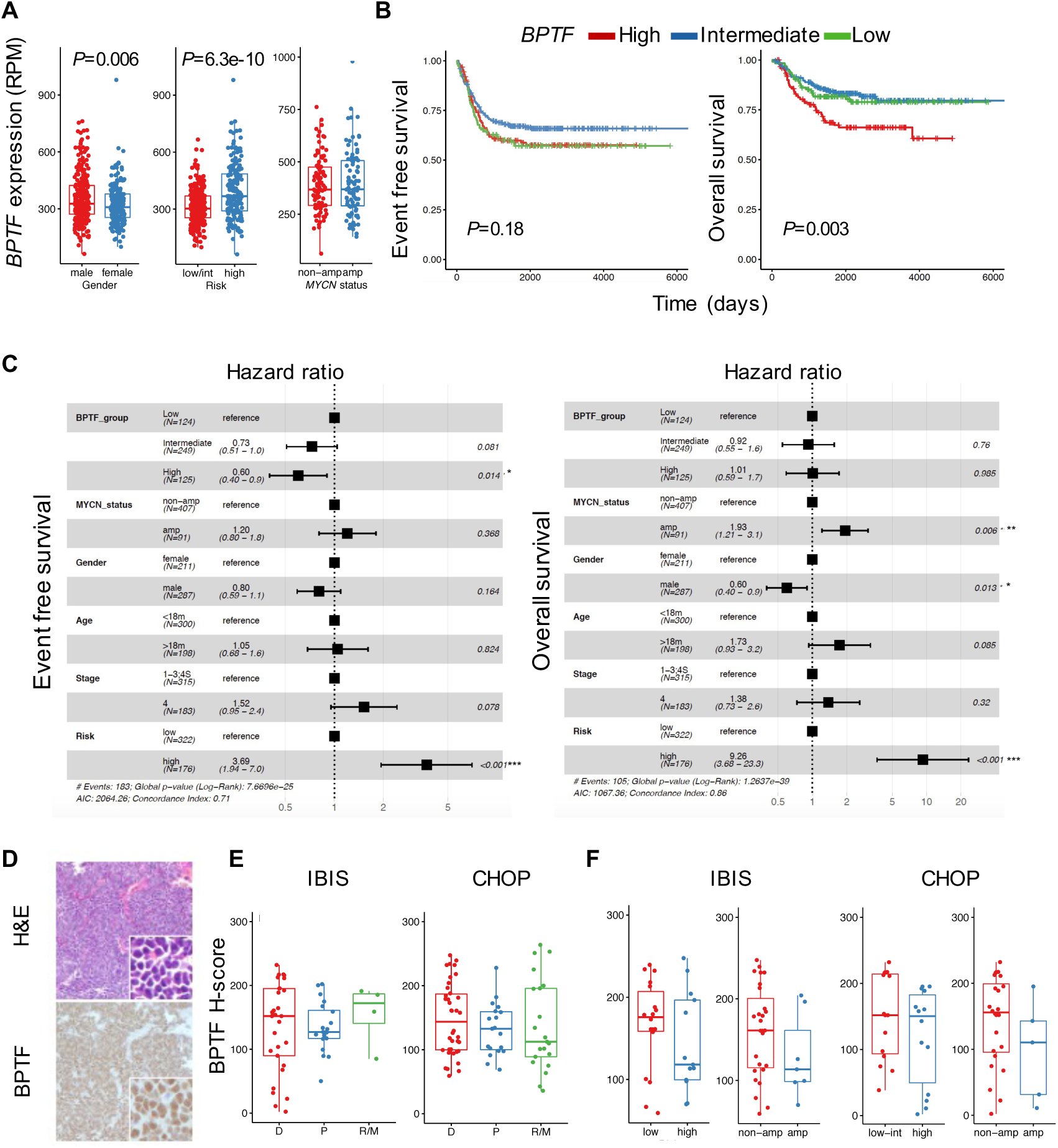
High *BPTF* expression in neuroblastoma samples is associated with high-risk and outcome. (A) *BPTF* is expressed at higher levels in primary neuroblastoma tumors from males and in tumors from high-risk patients (SEQC RNA-seq dataset, n=498). In contrast, expression is not associated with *MYCN* status in samples from high-risk patients (n=176) (Wilcoxon test). (B) High *BPTF* expression is associated with significantly worse overall survival (High, Q1 intermediate, Q2+Q3 low, Q4) (Log-rank test). (C) Cox proportional hazards regression models showing the association of the main clinical covariates and BPTF expression groups with EFS and OS in the SEQC dataset. Error bars represent the 95% confidence intervals and the indicated p values are from log-rank test. Sample size of each group is indicated in the figures. (D) BPTF protein is highly expressed in human neuroblastoma samples. A representative image of H&E and BPTF staining is shown. (E) BPTF protein expression levels are not associated with time of tumor sampling (D=diagnosis, P=post-treatment, or R/M=relapse/metastasis) in two independent series [IBIS (D=38, P=19, R/M=21) (left) and CHOP (D=32, P=22, R/M=4) (right)]. (F) BPTF expression is similar regardless of risk group or *MYCN* status in primary tumors of two independent patient series (IBIS and CHOP).

We also explored whether expression was associated with other genetic features of neuroblastoma in the Therapeutically Applicable Research to Generate Effective Treatments (TARGET) cohort, consisting mainly of high-risk tumors, and in the GMKF cohort. We found a positive correlation between *BPTF* mRNA levels and the read coverage of chr 17q, a surrogate of gene copy number, in both cohorts (TARGET, R=0.57, *P*=3.1e-10); GMKF, R=0.73, *P*<2.2e-16) (**Figure S3A**). Other genomic alterations relevant to neuroblastoma - such as chr 1p and 11q loss and *ALK* mutations - were not associated with *BPTF* expression levels in the TARGET dataset **(Figure S3B**). To determine whether methylation might contribute to determine *BPTF* expression levels, we assessed the distribution of beta-values at CpG dinucleotides in the *BPTF* locus. We found variability in the distribution of beta-values in the *BPTF* promoter; intragenic methylation was weakly correlated with *BPTF* expression levels (**Figure S3 C-E**). These results strongly suggest that 17q copy gain is a main determinant of *BPTF* expression levels.

To confirm BPTF expression in patient tumors, we used IHC on tissue microarrays (TMAs). We analyzed three cohorts, two of which comprised human tumor samples obtained at diagnosis, post-treatment, or at relapse/metastasis (**Figures 2D, E and S4**). High BPTF histoscores were obtained in both series. There was no association between BPTF expression levels and time of biopsy (**Figure 2E**). In primary tumors, there was no association of histoscores with gender, age, risk group, or *MYCN* amplification (**Figure 2F and S4**). Similar findings were made in a third series of samples from patient-derived xenografts (PDX) (**Figure S4**).

In conclusion, BPTF is overexpressed across neuroblastomas and higher BPTF expression is associated with high-risk neuroblastoma, but it is not an independent prognostic factor.

**Figure S3.**
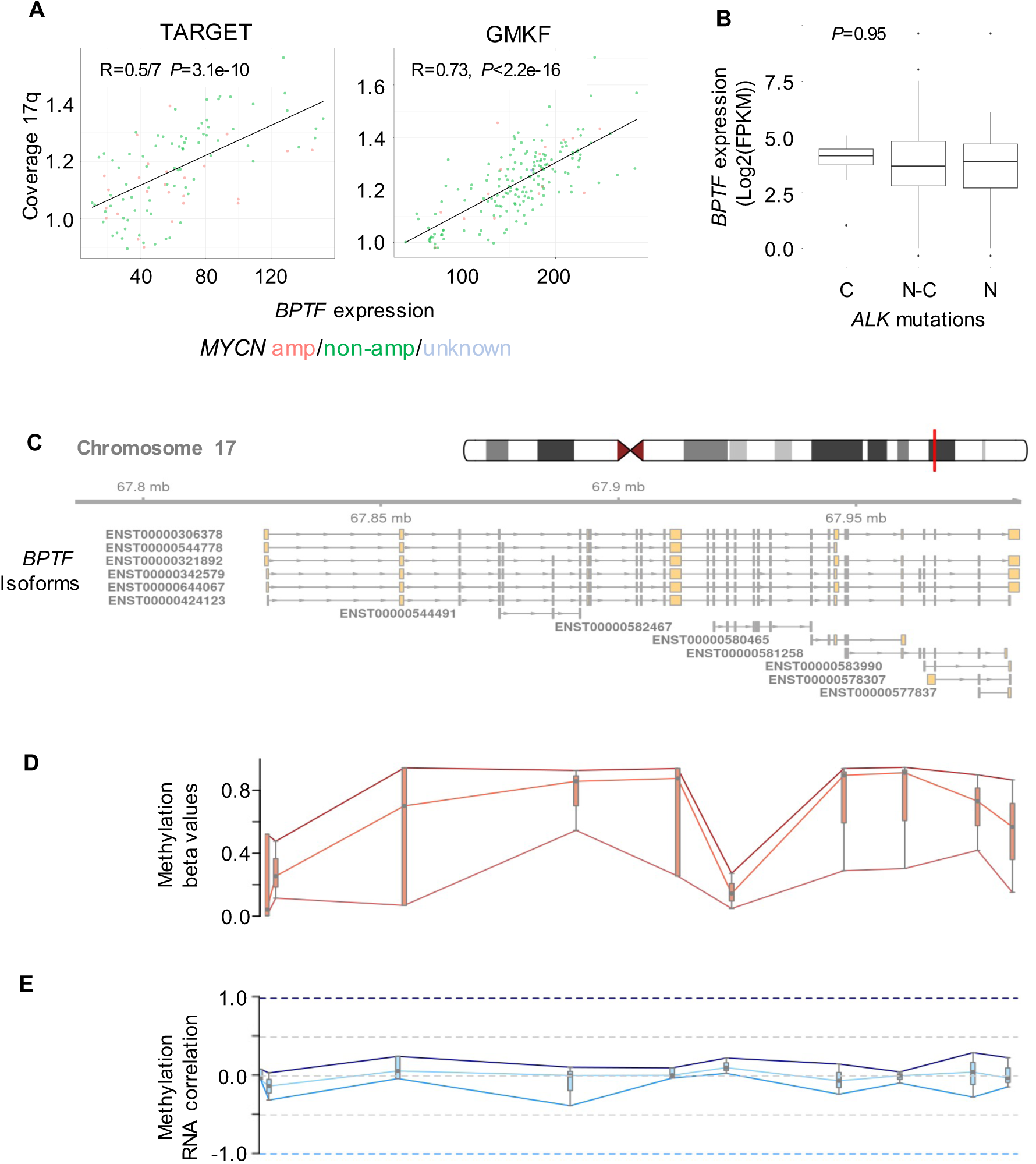
Correlation of genomic alterations and methylation with *BPTF* expression. (A) *BPTF* mRNA expression levels in patient tumors correlates directly with chr17q read coverage in the TARGET and GMKF cohorts (Pearson, R). (B) *BPTF* expression across patients with canonical (C, F1174L or R1275Q), non-canonical (N-C), and novel (N) *ALK* mutations in the TARGET dataset (N=105) (ANOVA test). (C) *BPTF m*RNA expression levels correlate weakly with DNA methylation at the *BPTF* locus in the TARGET dataset (n=115); *BPTF* mRNA isoforms are shown. (D) Distribution of beta-values at CpG dinucleotides in the *BPTF* locus, with the red lines tracing the maximum, median, and minimum beta-values at each position. (E) Boxplots showing the distribution of Pearson correlation coefficients of beta-values with the mRNA expression levels of each *BPTF* isoform. Blue lines trace the maximum, median, and minimum correlations at each location. The beta-value statistic ranges from 0-1; ‘0’ indicates all copies of the CpG dinucleotide in each sample are completely unmethylated (hypomethylation) and a value of ‘1’ indicates that every copy is methylated (hypermethylation) ^31^.

**Figure S4.**
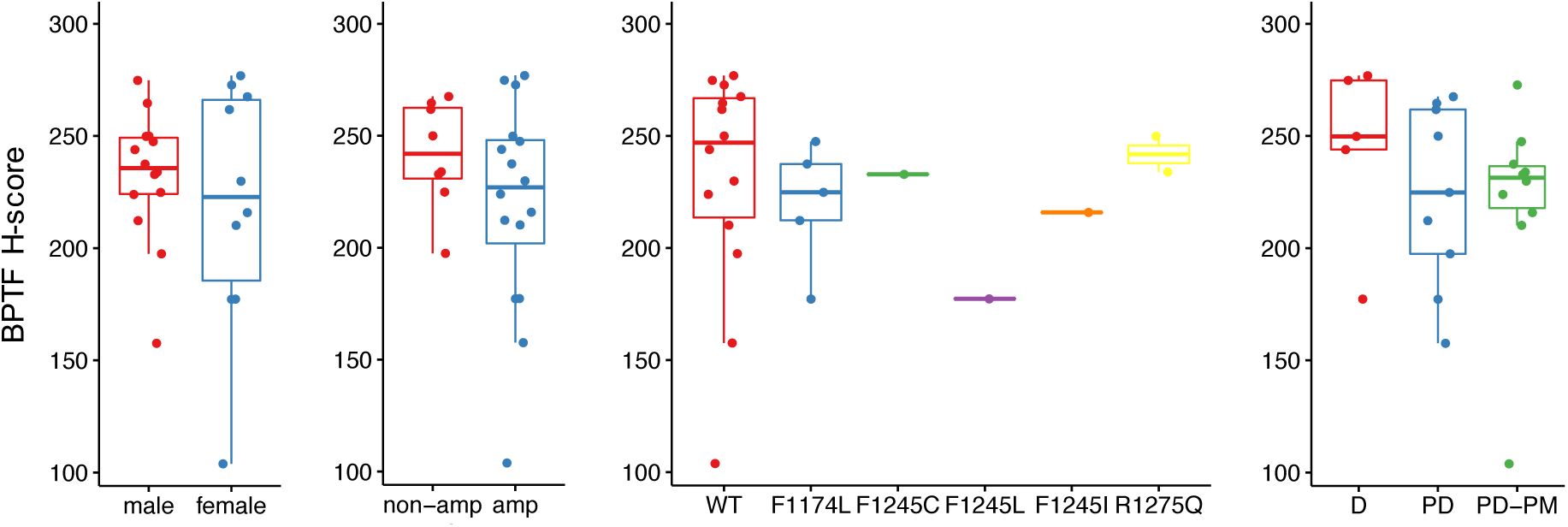
*BPTF* expression in neuroblastoma PDXs arrayed as TMA. Analysis of BPTF expression in the PDX CHOP-TMA and the association with sex, *MYCN* status, *ALK* mutation and phase of therapy.

### *BPTF* expression levels are associated with high cell cycle activity

To identify genetic programs linked to *BPTF* overexpression, we compared the transcriptome of neuroblastomas with the highest (top 10%) vs. the lowest (bottom 10%) *BPTF* mRNA levels in the SEQC dataset. In total, 2871 genes were significantly overexpressed (log_2_FC>2, adj. *P*<0.05) and 4314 genes were expressed at lower levels (log_2_FC<-2, adj. *P*<0.05) in *BPTF*-high tumors. GSEA revealed that *BPTF*-high tumors were significantly enriched in cell cycle-related pathways (e.g., G2/M checkpoint, mitotic spindle, and E2F targets) (*P<*0.05, FDR<0.25). *BPTF*-low samples were significantly enriched in epithelial-mesenchymal transition (EMT), inflammation, and NF-κB pathways, among others (**Figure 3A**). These findings were confirmed when using the same approach with the HR-TARGET dataset. Promoter scanning analysis of the overexpressed genes showed significant enrichment of the NFY, E2F, GATA, MYC, and SOX binding motifs whereas promoters of genes expressed at lower levels were enriched in the IRF1 and RELA motifs (**Figure 3B**). Overall, these results indicate that *BPTF* expression is associated with a highly proliferative neuroblastoma cellular state.

**Figure 3.**
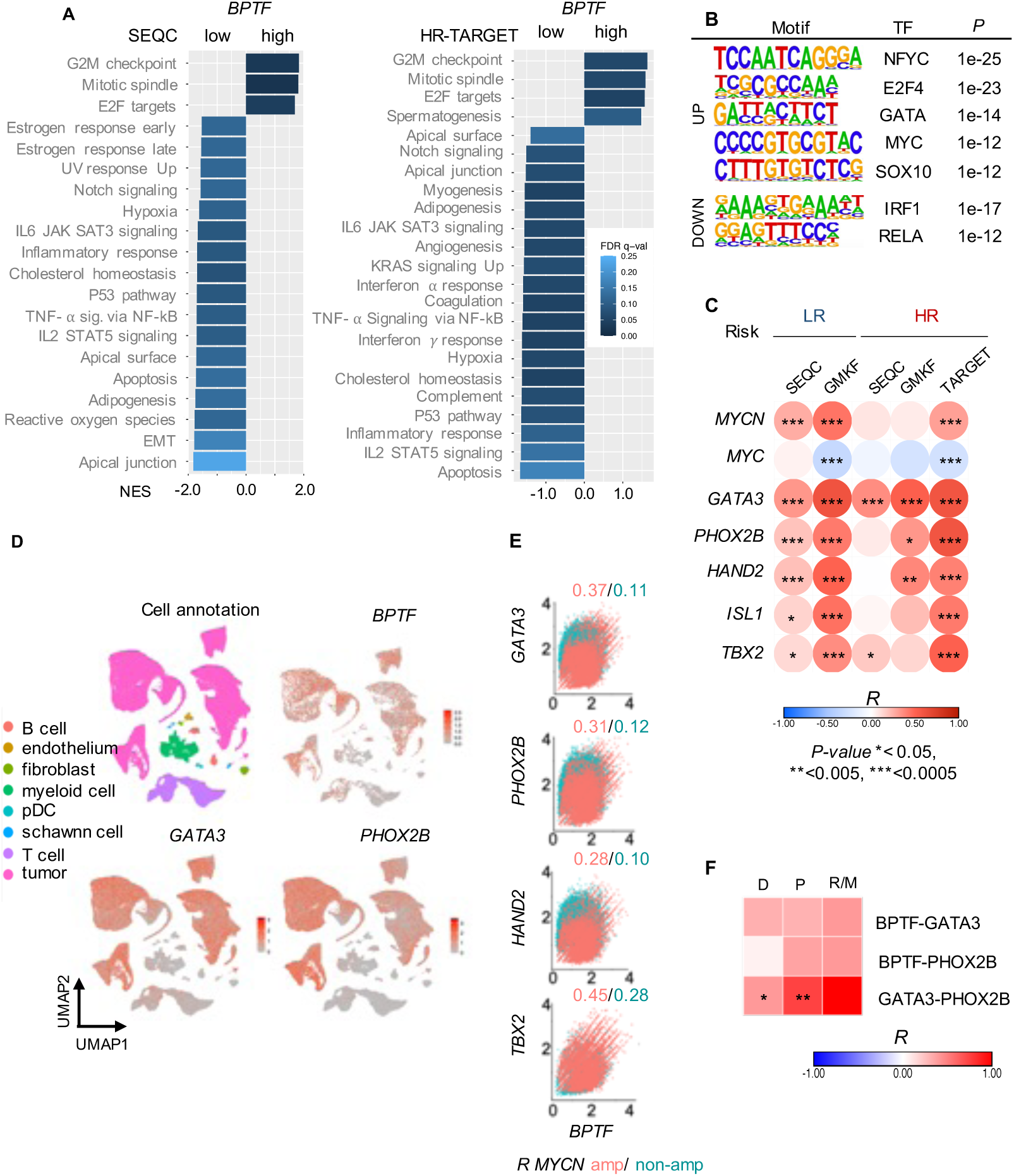
*BPTF* expression in neuroblastoma patient samples correlates with expression of the CRC transcription factors and with cell cycle related transcriptomic signatures; *BPTF* is mainly expressed in tumor cells. (A) Pathway enrichment analysis comparing the highest (1st decil) vs. the lowest (last decil) *BPTF* mRNA level subgroups in the SEQC (left) and HR-TARGET (right) datasets, using GSEA (nom P-val<0.05, FDR>0.25). (B) Main motifs associated to the enriched pathways identified through promoter scanning analysis of over- and under-expressed genes in the SEQC dataset (HOMER). (C) Expression of *BPTF* and the CRC transcription factors is strongly and directly correlated in the complete SEQC and HR-TARGET RNA-seq datasets (Spearman). (D) Single cell RNA-Seq analysis shows that *BPTF* expression is mainly restricted to neuroblastoma tumor cells ^32^. *GATA3* and *PHOX2B* are shown as representative members of the neuroblastoma CRC. (E) Correlation of the expression of *BPTF* and CRC components at the single cell level in *MYCN-*amplified vs non-amplified cells (R value, Pearson, P-val <2.2e-16). (F) Heatmap representing the correlation of the H-scores for BPTF, GATA3, and PHOX2B in the CHOP TMA series stratified according to the time of sampling (D, Diagnosis P, Post-treatment R/M=Relapse/Metastasis).

### *BPTF* and CRC component expression levels are positively correlated

The enrichment of the GATA and MYC motifs in genes up-regulated in *BPTF*-high tumors points to a functional interaction between BPTF and the transcription factors of the ADRN neuroblastoma CRC. To address this hypothesis, we mined the patient RNA-seq datasets (**Figure 3C**). Expression levels of *BPTF* and the CRC components were strongly and positively correlated both in low- and high-risk subgroups. The highest correlation was found with *GATA3* (Pearson, R=0.59, *P*=4.70E-013), *PHOX2B* (R=0.59, *P=*4.50E-13), and *TBX2* (R=0.51, *P*=1.00E-09) in the TARGET dataset. *BPTF* expression levels were significantly and positively correlated with *MYCN* expression in the SEQC cohort (R=0.27, *P*=1.2E10-9), the strength of the correlation being somewhat lower in high-risk tumors of the TARGET dataset (R=0.3, *P*=6E10-4). In contrast, there was no correlation with *MYC* expression in the SEQC cohort. To confirm and extend these findings, we analyzed the single cell RNA-seq data from Dong et al, 2020 ^32^, including 15 tumors and 144571 cells, 76% of which were malignant. *BPTF* was expressed at highest levels in neuroblastoma and T cells (**Figure 3D**). *GATA3* and *PHOX2B* showed a similar distribution in tumor cells. At the single cell level, a positive correlation of *BPTF* expression with CRC members was present in tumor cells (**Figure 3E**). Pearson coefficient was estimated separately for *MYCN* amplified/non-amplified tumor cells, being higher in *MYCN*-amplified cells, especially with *GATA3* (R=0.37, *P*<2.2e-16) and *TBX2* (R=0.45, *P*<2.2e-16) (**Figure 3E**).

The RNA-based correlations described above were largely confirmed at the protein level using TMAs immunostained for BPTF, GATA3, and PHOX2B. There was a modest positive correlation of BPTF with GATA3 (R=0.33, *P*= 0.054) in the PDXs series and in patient tumors, although the findings were only significant for GATA3-PHOX2B expression in diagnostic and post treatment samples (R=0.4, *P*=0.028; R=0.74, *P*=3e-4, respectively) (**Figure 3F**). The lower statistical significance at the protein level may be related to the smaller sample size.

These findings support a functional interaction between BPTF and the neuroblastoma CRC.

### BPTF and CRC transcription factors are part of the same molecular complex

To investigate the functional interplay observed between BPTF and the CRC components, we first compared their subcellular distribution using immunofluorescence. As expected, BPTF, GATA3, PHOX2B, and HAND2 co-localized in the nucleus of ADRN neuroblastoma cells (**Figure 4A**). Immunoprecipitation (IP) followed by WB revealed that BPTF and GATA3 are present in the same complex in KELLY and LAN-1 cells (**Figures 4B and S5B**). However, we could not detect the other CRC components in BPTF immunoprecipitates. To acquire more resolution, we analyzed BPTF immunoprecipitates from neuroblastoma cell lysates by mass spectrometry (**Figures 4C-E and S5C**). Two-hundred-and-five and 75 significant interactors were detected in KELLY and in SK-N-AS cells, respectively. BPTF was amongst the most significantly enriched proteins in both cell lines (KELLY, log_2_FC BPTF/IgG=6.38, FDR=0.066) and (SK-N-AS; log2FC BPTF/IgG=5.76, FDR=0.066). Several NURF complex components - such as BAP18 (C17orf149), SMARCA proteins, and RBBP4 - were identified in immunoprecipitates from both cell lines (**Figure 4D**), thus validating the experimental strategy. Interactors network K-means clustering showed BPTF as a member of the chromatin remodeling and histone modification cluster (**Supplementary Table 2**). Twenty-three proteins were identified in immunoprecipitates from both lines (**Figure S5D**), with DAB2IP and IQGPA1 being the top ones. Several of these interactions were validated by co-IP/WB (**Figure S5E**). MYCN and HAND2 were found in KELLY immunoprecipitates, further supporting that BPTF binds the CRC (**Figure 4C**). A direct BPTF-MYCN interaction was confirmed by PLA in LAN-1 (**Figure S5F**). Besides MYCN and HAND2, other proteins relevant to the ADRN program were identified, including MAX, ELAVL4, and DBH. Pathway analysis of BPTF interactors showed enrichment in cell cycle-related proteins only in KELLY cells (**Figure 4E**); in contrast, interferon response proteins were enriched in SK-N-AS cells (**Figure S5G**). Altogether, these results support a physical and functional interaction of BPTF and the ADRN CRC and suggest distinct partnerships associated with cell lineage identity.

**Figure 4.**
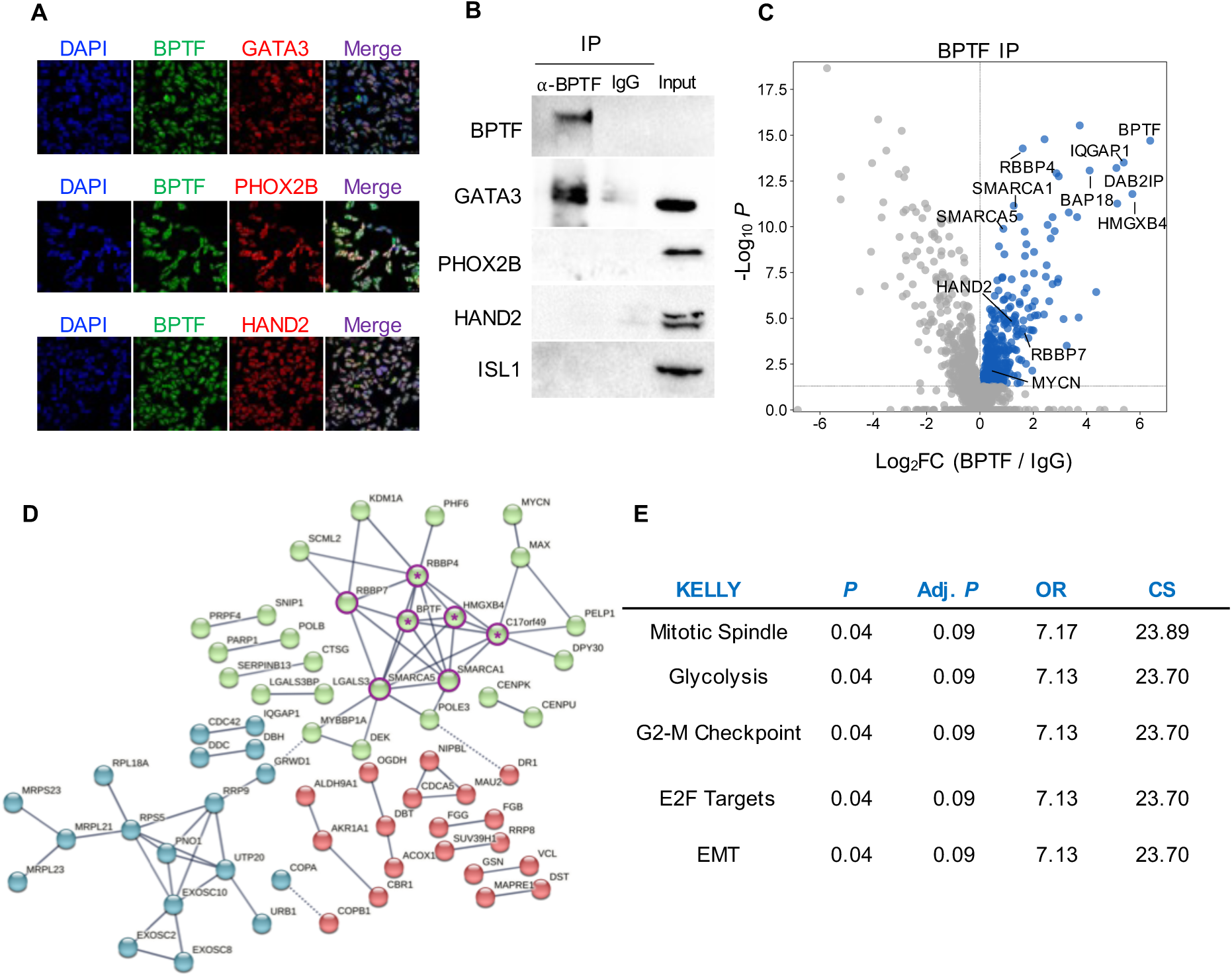
*BPTF* and transcription factors of the ADRN CRC co-localize in neuroblastoma cells and are present in the same complex in KELLY cells. (A) Nuclear colocalization of BPTF and the CRC factors GATA3 and HAND2. (B) Co-IP and WP show that BPTF and GATA3 are present in the same complex. (C) Mass-spectrometry analysis of BPTF immunoprecipitates identifies HAND2 and MYCN as BPTF interactors. (D) STRING interaction network derived from the IP-MS experimental data (interaction score=0.9.) The edges indicate both functional and physical protein associations; line thickness refers to the strength of data (K-means clustering for 3 clusters (Supplementary Table 2); edges between clusters indicated by dotted line; purple circle, NURF component; *, also found in the SK-N-AS IP-MS experiment). (E) Pathway enrichment analysis of BPTF significant interactors (Enrichr). OR, Odds ratio; CS, Combined Score.

**Figure S5.**
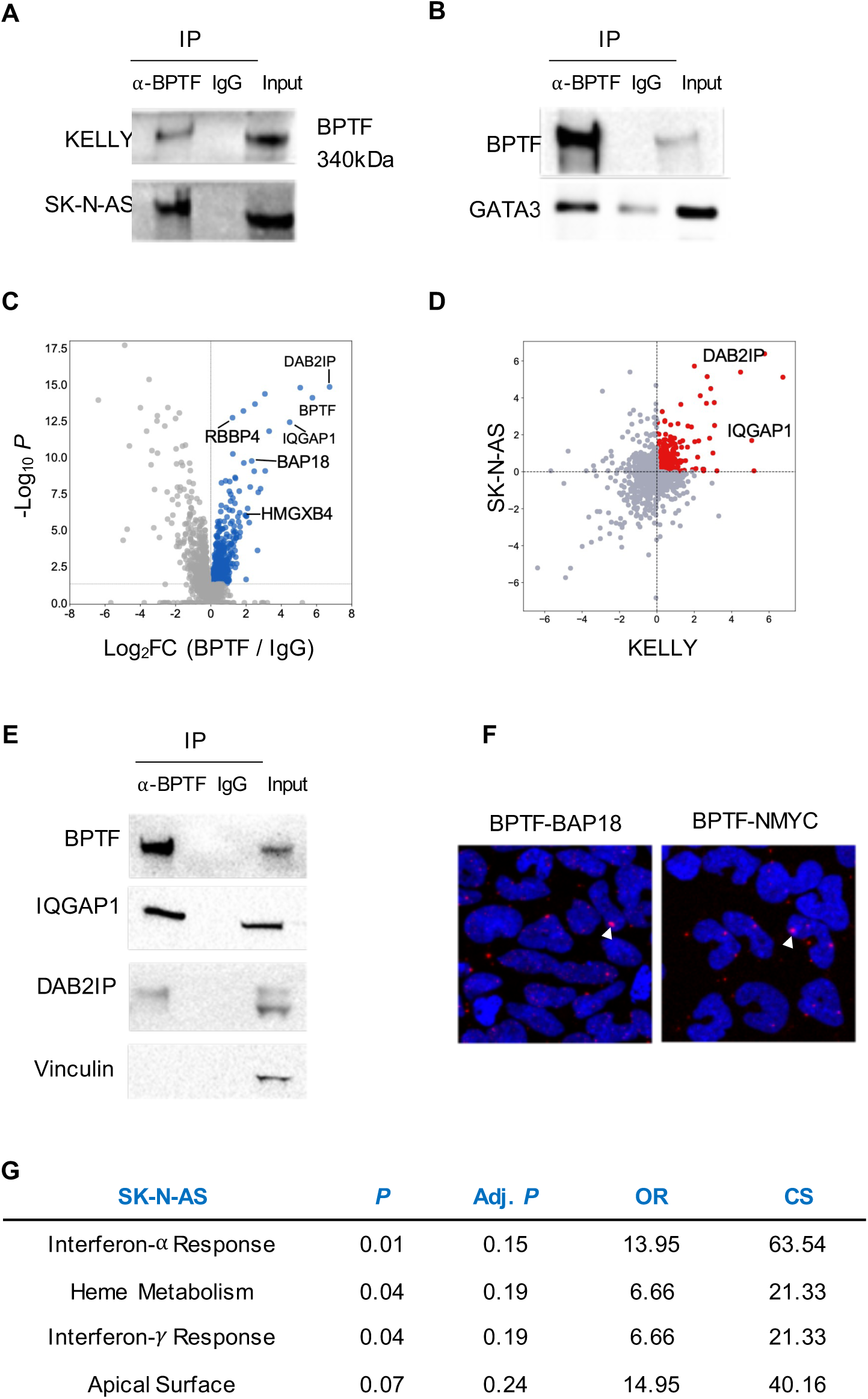
BPTF interactors detected by IP/MS in neuroblastoma cells. (A) Validation of the BPTF IP by WB using KELLY and SK-N-AS cells. One representative image of 3 independent biological replicates is shown. (B) Validation of the BPTF-GATA3 interaction in LAN-1 cells using co-IP-WB. (C) Volcano plot showing with red dots the significant interactors detected by IP/MS in SK-N-AS cells. (D) Graph showing the common significant BPTF interactors in KELLY and SK-N-AS cells. (E) Co-IP-WB validation of selected top common interactors using KELLY cells. (F) PLA for BPTF-BAP18, a known BPTF interactor, and BPTF-MYCN in LAN-1 cells. The interaction events are visible as red dots (nuclear staining in blue) (See white arrows). (G) Representative significantly enriched pathways from the list of BPTF interactors in SK-N-AS cells by Enrichr. OR=Odds ratio, CS=Combined Score.

**Supplementary Table 2.**
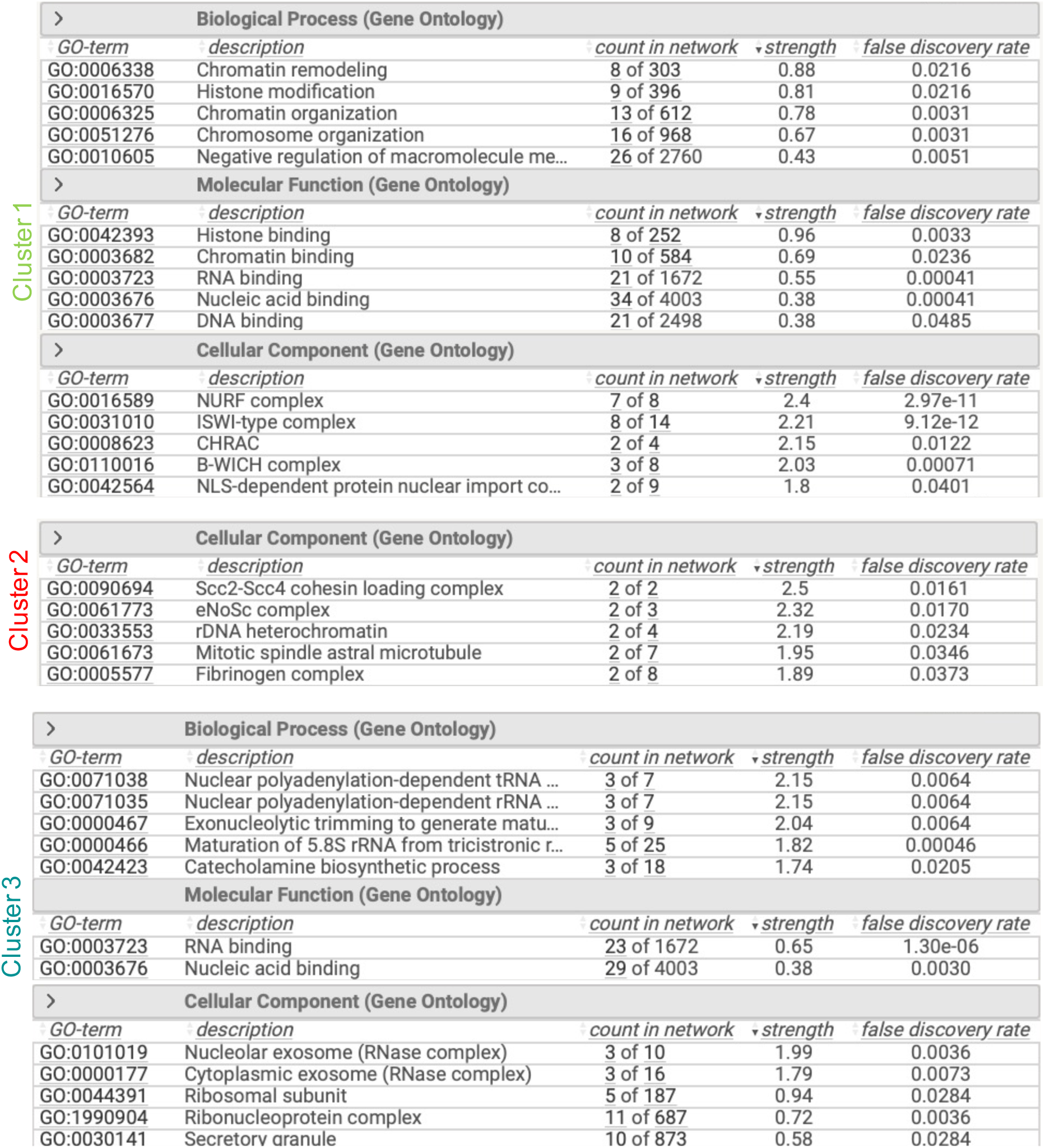
Top significant Gene Ontologies from K-means clustering of BPTF interactors in KELLY cells.

### MYCN and MYC bind to the *BPTF* promoter and regulate its expression

To assess the functional relationship between BPTF and MYCN/MYC we have leveraged on the extensive information on the genomics of neuroblastoma cell lines. In a panel of 17 neuroblastoma lines, mRNA expression of *BPTF* and CRC components was similarly high across all lines (**Figure 5A**). In contrast, *MYCN* and *MYC* transcript levels were high in ADRN and MES lines, respectively, and were negatively correlated (**Figure 5A**, Spearman R= -0.57, *P*=0.017). We have chosen KELLY, LAN-1, and KP-N-YN as prototypic ADRN lines, and SK-N-AS as the most MES line for further experiments. To determine whether MYC proteins and the neuroblastoma CRC are involved in the regulation of *BPTF* expression, we leveraged public ChIP-Seq data of MYCN and MYC in *MYCN-*amplified (COGN415, KELLY, LAN-5, NGP, and NB1643) and *MYCN*-non-amplified (NB69, SK-N-AS, and SK-N-SH) neuroblastoma cells^33^. **Figure S6** shows MYCN binding to *BPTF* in all *MYCN*-amplified cell lines tested (n=5). Similarly, MYC was found to bind to *BPTF* in the *MYCN*-non-amplified cells (n=3). The *LIN28A* region is shown as binding control (**Figure S6A**, right panel). Focusing on the *BPTF* promoter, MYCN binds to this region in 5/6 *MYCN*-amplified cell lines, although with variable enrichment, the signal being highest in KELLY and NB1643 cells (**Figure S6B**). MYC binding to the *BPTF* promoter was comparatively weaker (**Figure S6B**). We could not find evidence for the ADRN CRC proteins binding to the *BPTF* promoter (-3Kb to +1.5Kb from the TSS), distal enhancer, or super-enhancer (SE) regions, defined from a dataset resulting from the merge of regions identified in cultured cells and patient tumors (NB-SE) (n=2800) ^25, 34^, except that a PHOX2B peak was found in the *BPTF* distal enhancer (**Figure S6C**).

**Figure 5.**
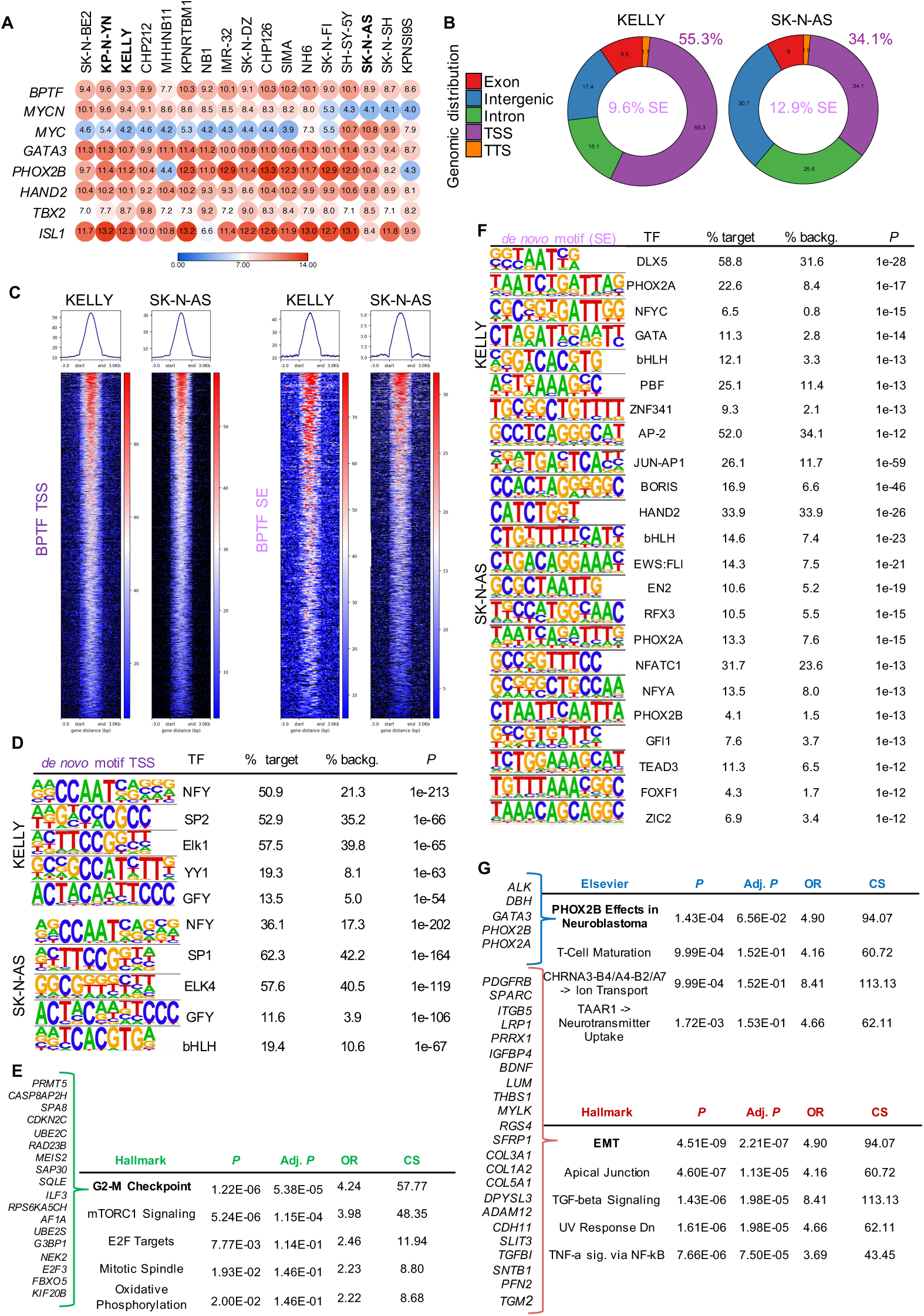
BPTF binds preferentially to TSS regions of genes related to cell cycle control. (A) Heatmap showing the mRNA expression of *BPTF*, *MYCN*, *MYC,* and CRC factors in neuroblastoma cell lines ordered by decreasing *MYCN* expression levels. (B) Genomic distribution of the overlap of annotated peaks in Cut&Run experiments using KELLY (≥3 replicates) and SK-N-AS (≥2 replicates) cells. (C) Heatmap showing the binding of BPTF to the TSS and SE regions in a representative experiment with both cell lines. (D) *De novo* top 5 motifs in the promoter-TSS in KELLY and SK-N-AS cells. (E) Pathway analysis of common annotated genes in promoter-TSS in the 4 cell lines used (KELLY, LAN-1, KP-N-YN, SK-N-AS). The top 5 pathways are shown. The main genes involved in the top pathway are listed. (F) *De novo* significant top motifs in SE regions in KELLY and SK-N-AS cells. (G) Pathway enrichment from the overlap of the annotated genes in SE regions in ≥2 ADRN cell lines (top, blue) and the specific annotated genes for the MES cell line SK-N-AS (bottom, red). The main genes involved in the top pathway are listed.

### BPTF cooperates with MYC proteins at TSS regions to regulate cell cycle-related genes

To better understand the functional interaction between BPTF, MYCN, and CRC components, we analyzed whether BPTF regulates the latter. Little is known about the genome-wide distribution of BPTF, due to technical limitations. We performed ChIP-Seq and Cut&Run experiments in three ADRN (KELLY, LAN-1, and KP-N-YN) and one MES (SK-N-AS) neuroblastoma cell lines with multiple antibodies, but only Cut&Run provided robust data on the genomic distribution of BPTF. Peaks with signal value >3 and a significant FDR <0.05 were annotated using HOMER and the binding was summarized considering the overlap of peaks identified in at least 3 (out of 5) biological replicates in KELLY and in at least 2 (out of 4, 4, and 3 biological replicates, respectively) in the other cells. In all cases, BPTF peaks mapped mainly at TSS-promoter regions, especially in KELLY (55.3% of all peaks) and LAN-1 (58.3%) cells (**Figures 5B and S7A**). In addition, 6.9-13.4% of BPTF peaks mapped to SE regions (**Figures 5B, C and S7A**). We then examined the overlap of BPTF binding regions with public ATAC-seq data ^34^: 97.2% and 94.3% of promoter/TSS-annotated peaks were in open chromatin regions in KELLY and in SK-N-AS cells, respectively (**Figure S7B**).

**Figure S6.**
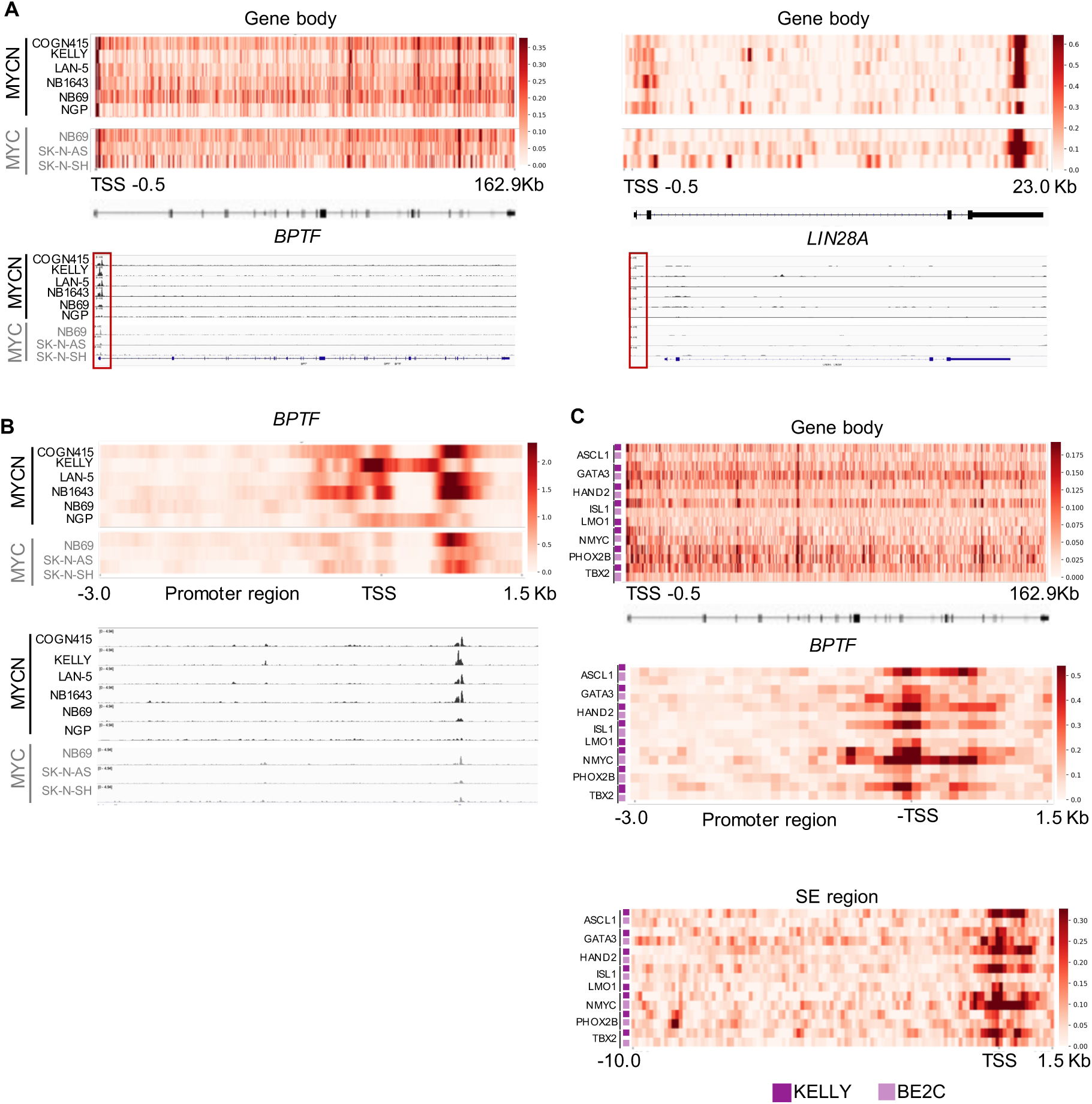
ChIP-seq analysis of the distribution of CRC transcription factors at the *BPTF* locus and regulatory regions in neuroblastoma cells. (A) Heatmap of MYCN-ChIP and MYC-ChIP signal in *BPTF* (left) and *LIN28A* genes (entire gene region) (right) (Top). IGV screenshots of MYCN and MYC binding to the *BPTF* and *LIN28A* gene body (bottom). The *LIN28A* region is shown as a representative gene without MYCN/MYC signal enrichment. (B) Heatmap of MYCN-ChIP and MYC-ChIP signal in the *BPTF* promoter region (-3.0Kb to +1.5Kb from TSS). IGV screenshots of binding of MYCN and MYC to the *BPTF* promoter. (C) Heatmap of CRC-ChIP signal in the *BPTF* gene (entire gene), promoter (-3.0Kb to +1.5Kb from TSS) and SE region (10kb upstream of the TSS).

**Figure S7.**
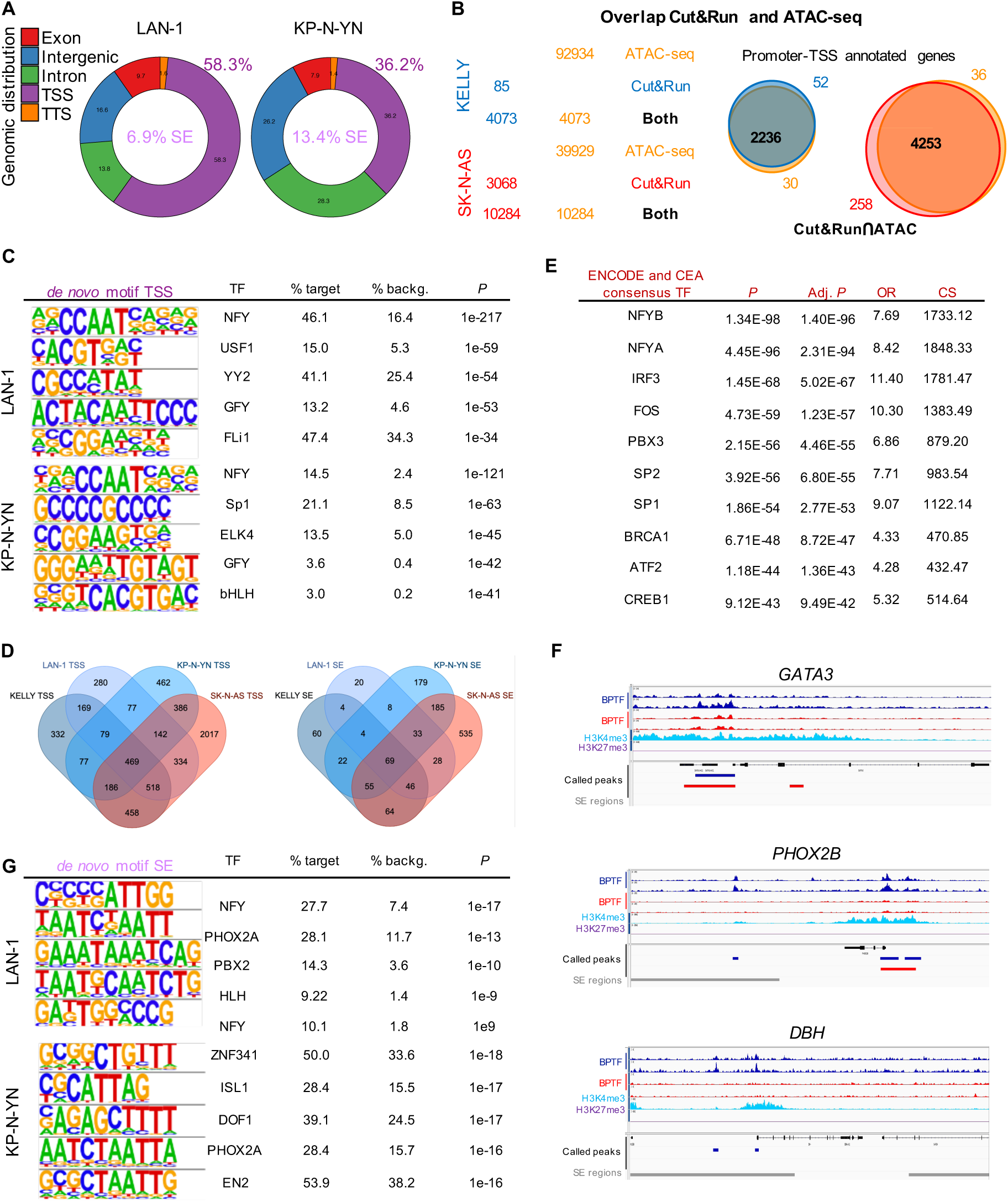
Analysis of the genome wide distribution of BPTF assessed using Cut&Run. (A) Genomic distribution of the overlap of annotated peaks in Cut&Run experiments using LAN-1 and KP-N-YN cells (≥2 replicates). (B) Overlap of the Cut&Run peaks with ATAC-seq open regions in KELLY (blue) and SK-N-AS (red) cells. Venn diagram of the overlap of annotated genes in TSS regions in the Cut&Run experiment with the accessible chromatin regions. (C) *De novo* top 5 transcription factor binding motifs in the TSS-annotated peaks (HOMER) from the BPTF Cut&Run experiments. (D) Venn diagrams showing the overlap of annotated genes in TSS-promoters (left) and SE (right) in KELLY, LAN-1, KP-N-YN, and SK-N-AS cells. (E) Transcription factor binding motif analysis of common annotated genes in promoter-TSS regions in the 4 cell lines (Enrichr). (F) Representative IGV screenshots of BPTF binding to *GATA*3 (top), *PHOX2B* (middle), and *DBH* (bottom) in KELLY (blue) and SK-N-AS (red) cells. H3K4me3 and H3K27me3 peaks are shown for comparison. (G) Transcription factor binding motif analysis of SE-annotated peaks (HOMER) from the BPTF Cut&Run experiments.

To gain further insights into the role of BPTF in transcriptional regulation, we performed *de novo* motif analysis of the BPTF TSS-mapping reads in KELLY cells using HOMER. This revealed an enrichment of NFY, SP2, ELK1, YY1, and GFY binding sequences (**Figure 5D**). Similar findings were made in the other three lines, NFY being the top motif identified in all of them (**Figure 5D and S7C**). Remarkably, SP1/2 motifs were also found among the top identified. The overlap of TSS-annotated genes in the four lines consisted of 469 genes associated with NFY transcription factors and related to late cell cycle pathways and mTORC1 signaling (**Figure 5E and S7D, E**). Similar pathways emerged in ADRN and MES cells, when analyzed separately. Regarding the CRC components, BPTF binding was found at the promoter of *PHOX2B* and *HAND2* in all the cell lines analyzed and at the promoter of *GATA3*, *TBX2,* and I*SL1* in 3 out of 4 lines; representative peaks are shown in **Figure S7F**. A BPTF peak was also found at the promoters of *MYCN* and *MYC* in LAN-1 and SK-N-AS cells, respectively.

To assess the functional effects of *BPTF* loss-of-function on the CRC factors, we used an inducible lentiviral *BPTF* knockdown in KELLY cells (**Figure S8A**) and performed RNA-seq at 48h (**Figure S8B**): 163 and 14 genes were significantly down-regulated and up-regulated, respectively. *BPTF* knockdown resulted in significant reduction of *MYCN* mRNA expression (log2FC -0.90, P-adj 0.006, **Figure S8B, C**). Consistent with the genomic binding data, *PHOX2A* was also significantly downregulated (log2FC=-0.87, adj *P*=0.007). Expression of *HAND2*, and *ISL1* was non-significantly reduced while *PHOX2B*, *GATA3*, and *TBX2* mRNA levels were similar in both conditions. Independent *BPTF* knockdown experiments using siRNAs showed reduced *PHOX2B* expression in KELLY cells (**Figure S8D**). Consistent with the results described above, the activity of MYC-related and mTOR signaling pathways was significantly down-regulated upon inducible *BPTF* knockdown (**Figure S8B**). Similarly, there was a down-regulation of the ADRN signature and - to a lesser extent - the MES signature ^25^ (**Figure S8E**).

To assess the effects of MYCN down-regulation, we treated KELLY cells with MYCi975 - a MYC binder a that disrupts MYC/MAX dimers and impairs MYC-driven gene expression ^35^- and found reduced expression of both MYCN and BPTF (**Figure S8F**).

Together with the reduced cell proliferation upon BPTF knockdown, these findings support the notion that BPTF regulates the expression of cell cycle genes through promoter binding and suggest that it regulates the activity of the CRC-driven programs.

**Figure S8.**
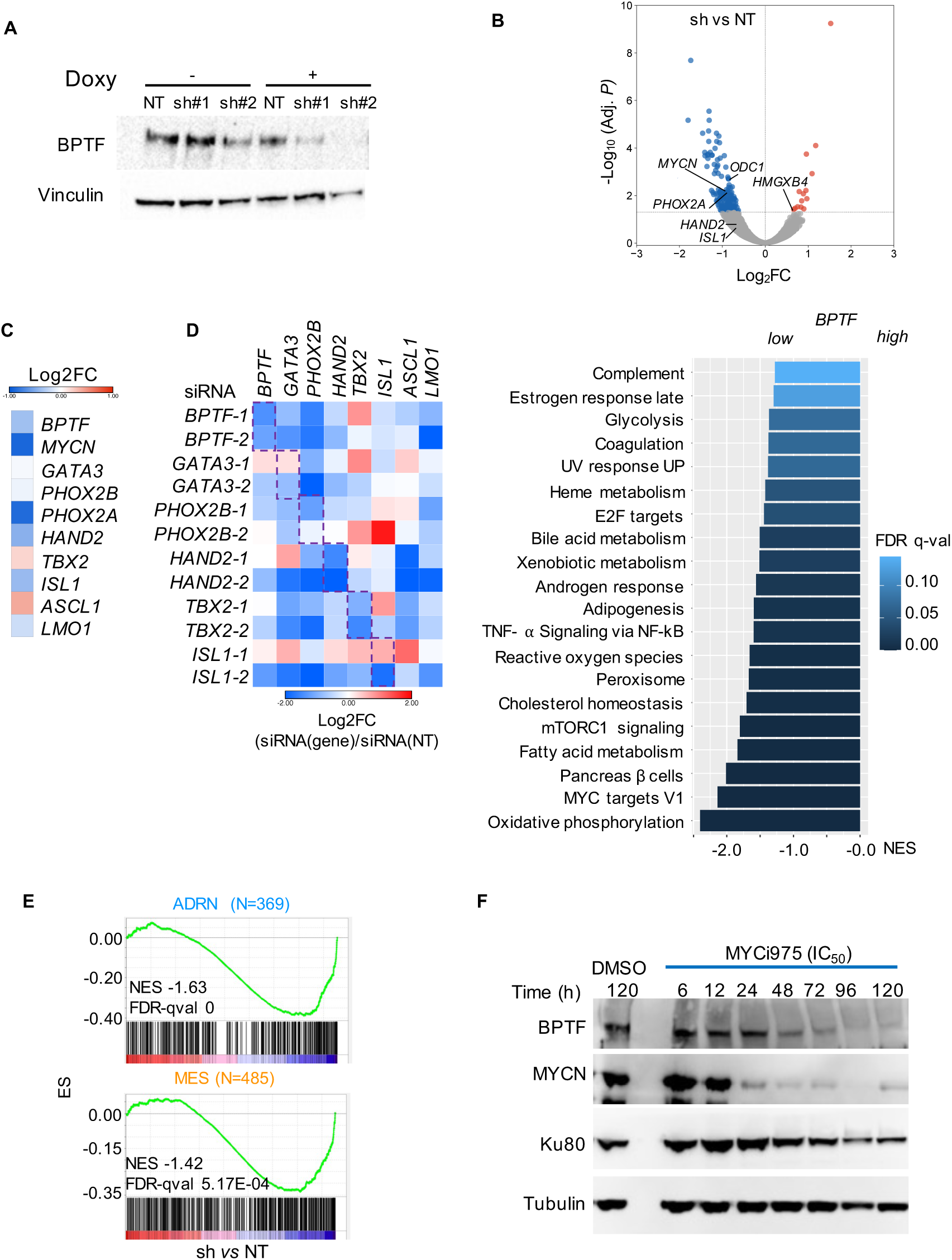
*BPTF* knockdown in KELLY cells leads to downregulation of CRC members and the cell identity signatures. (A) Reduction of BPTF protein levels 48h after induction of *BPTF* knockdown with doxycycline using two different shRNAs. (B) Volcano plot showing significant up- (red) and down-regulated (blue) genes upon inducible *BPTF* knockdown with sh2 (top). Significantly enriched pathways comparing sh2 vs NT (bottom). (C) Heatmap representing Log2FC expression of *BPTF* and CRC factors upon inducible *BPTF* knockdown. (D) Heatmap showing Log2FC (siRNA/NT) expression ratio for *BPTF* and CRC members 72h after knockdown with two different siRNAs targeting each gene. Dashed purple line highlights the gene downregulated with the corresponding siRNA. (E) Downregulation of the ADRN and MES signatures upon inducible *BPTF* knockdown. (F) Decreased expression of BPTF and MYCN after treatment of KELLY cells with MYCi975.

### BPTF binding to super-enhancers indicates cooperation with the neuroblastoma CRC

SE play a critical role in the control of neuroblastoma lineage identity. To better understand the role of BPTF in this process, we explored the intersection of BPTF Cut&Run peaks with the NB-SE dataset (see Methods) and found that the percentage of SE regions bound by BPTF ranged from 6.9-13.4%: 398 regions in KELLY, 246 in LAN-1, 698 in KP-N-YN, and 1722 in SK-N-AS cells. *De novo* motif analysis of peaks in SE regions revealed an enrichment in binding sequences for PHOX2A, NFY, GATA, and basic Helix-Loop-Helix (bHLH) proteins in ADRN cells (**Figures 5F and S7G**); the HAND2, bHLH, and PHOX2A/B motifs were also enriched in SK-N-AS cells, consistent with the fact that CRC components are expressed in these MES cells. Notably, other motifs related to EMT or MES subtypes - such as JUN, and TEAD - were enriched exclusively in SK-N-AS peaks. Among the SE-annotated genes in ≥2 ADRN cells were *GATA3*, *PHOX2A, PHOX2B*, *ALK*, and *DBH*, all of which are important for the ADRN subtype (**Figures 5G and S7F**). The genes annotated to the SE regions only in SK-N-AS cells were mainly associated with EMT, TGF-β signaling, and TNF-α signaling via NF-κB (**Figure 5G**), pathways related to a MES behavior. Among the included genes were *PRRX1*, several collagen family members, *PDGFRB*, and *TGFBI (***Figure 5G**).

We then assessed the co-occurrence of MYCN/MYC at the BPTF-bound promoters and/or SE regions (**Figure 6A, B**). Interestingly, MYCN co-localized with 95% of all BPTF peaks in KELLY cells and MYC co-localized with 68% of the BPTF peaks in SK-N-AS cells. When considering the TSS, the highest co-binding level was found for MYCN (KELLY) and MYC (SK-N-AS) (**Figure 6A, B**). The overlap with the remaining CRC factors at promoters was lower, ranging from 45% for TBX2 to 2.6% for ISL1. The proportion of genes co-bound at SE was similar for MYCN (97.7%) and MYC (66.5%). In contrast, the overlap with the CRC factors was higher at SE than at TSS regions, ranging from 77.4% for TBX2 to 50.5% for ISL1 (**Figure 6A**).

**Figure 6.**
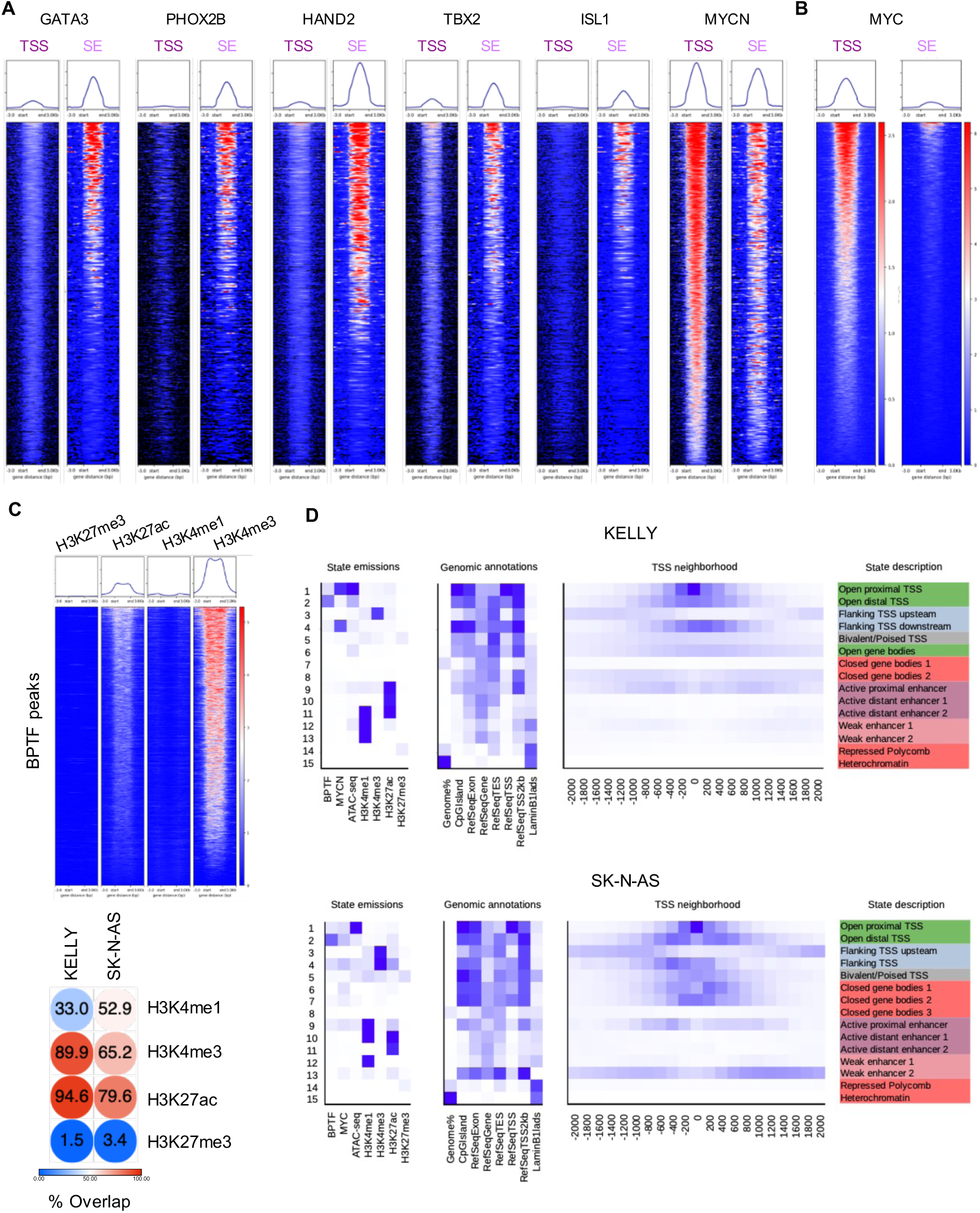
BPTF co-localizes at TSS regions with MYCN or MYC but shares SE positions with the CRC transcription factors. (A) Heatmap showing the intensity of the binding of MYCN and the CRC factors to TSS and SE regions where BPTF binds in KELLY cells. B) Heatmap showing the intensity of the binding of MYC to TSS and SE regions where BPTF binds in SK-N-AS cells. Representative scales for TSS and SE are shown. (C) Heatmap showing the intensity of the binding of the histone marks to BPTF peaks in KELLY cells and % of overlap of each histone mark with BPTF-bound regions in KELLY and SK-N-AS cells. (D) ChromHMM models summarizing the relative genome-wide distribution of BPTF and MYCN/MYC over chromatin states in KELLY (top) and SK-N-AS (bottom) cells. The panel on the left displays a heatmap of the emission parameters: each row corresponds to a different state and each column corresponds to a different mark (BPTF/Cut&Run; MCYN/MYC/ChIP-Seq; ATAC-seq; histone marks/ChIP-seq). The darker blue color corresponds to a greater probability of observing each mark in the corresponding state. The heatmap in the middle displays the overlap fold-enrichment for various external genomic annotations. The heatmap on the right shows the fold-enrichment for each state for each 200-bp bin position within 2 Kb around the TSS. A darker blue color corresponds to a greater fold-enrichment. State descriptions are based on ^36^.

We also assessed the co-localization of BPTF and histone marks (**Figure 6C**) using public ChIP-seq data for H3K4me3, H3K27Ac, H3K4me1 and H3K27me3 in KELLY and SK-N-AS cells ^33^. The percentage of H3K4me3 peaks coincident with BPTF peaks in KELLY and SK-N-AS cells was 89.9% and 65.2%, respectively. Accordingly, 94.6 and 79.6 % of H3K27Ac peaks overlapped with BPTF binding, respectively. A lower overlap was found with H3K4me1 and there was minimal overlap (1.5-3.4%) with H3K27me3. These findings strongly support that BPTF binding is largely restricted to active genes.

To address the hypothesis that the role of BPTF varies when partnering with MYCN or MYC, we compared the gene sets comprising BPTF-MYCN co-bound genes in KELLY and BPTF-MYC ones in SK-N-AS (**Figure S9**). The overlap of BPTF-MYCN with BPTF-MYC co-annotated genes was similar at TSS and SE (63% and 62%, respectively). Regarding the TSS, pathway analysis revealed that genes included in both overlapping gene sets were enriched in mTORC1 signalling, MYC-related, and G2-M checkpoint signatures; genes in the BPTF-MYC-only overlap were enriched in the E2F target pathway (**Figure S9A**). At SE, both BPTF-MYCN and BPTF-MYC overlapping genes were related to neuroblastoma cell identity (“PHOX2B effects in neuroblastoma”); genes in the BPTF-MYC-only overlap were also enriched in TGF-β, NF-κB signalling, and EMT pathways (**Figure S9B**). Altogether these results point to a shared role of BPTF with MYCN/MYC in TSS regions to regulate cell cycle but not at the SE level.

To assess the functional features of the genomic coordinates corresponding to BPTF and MYCN/MYC peaks, we used Poisson-based multivariate hidden Markov model (ChromHMM) ^36^. Chromatin state maps were generated using data from BPTF-Cut&Run, histone mark- and MYCN/MYC-ChIP-seq, and ATAC-seq data from KELLY and SK-N-AS cells (**Figure 6D**). The 15-state model was considered optimal based on exploratory analyses and the original description from Ernst and Kellis ^36^. BPTF signal was enriched in 4 chromatin states in KELLY and SK-N-AS cells: open proximal TSS, open distal TSS, flanking TSS downstream, and active proximal enhancers. In addition, the bivalent/poised TSS, closed gene bodies, and weak enhancers were also detected in SK-N-AS cells (**Figure 6D**) ^36^. MYCN/MYC binding shared these chromatin states. These results reinforce a major role of BPTF at TSS, its partnership with MYC family proteins, and an additional role at enhancers.

**Figure S9.**
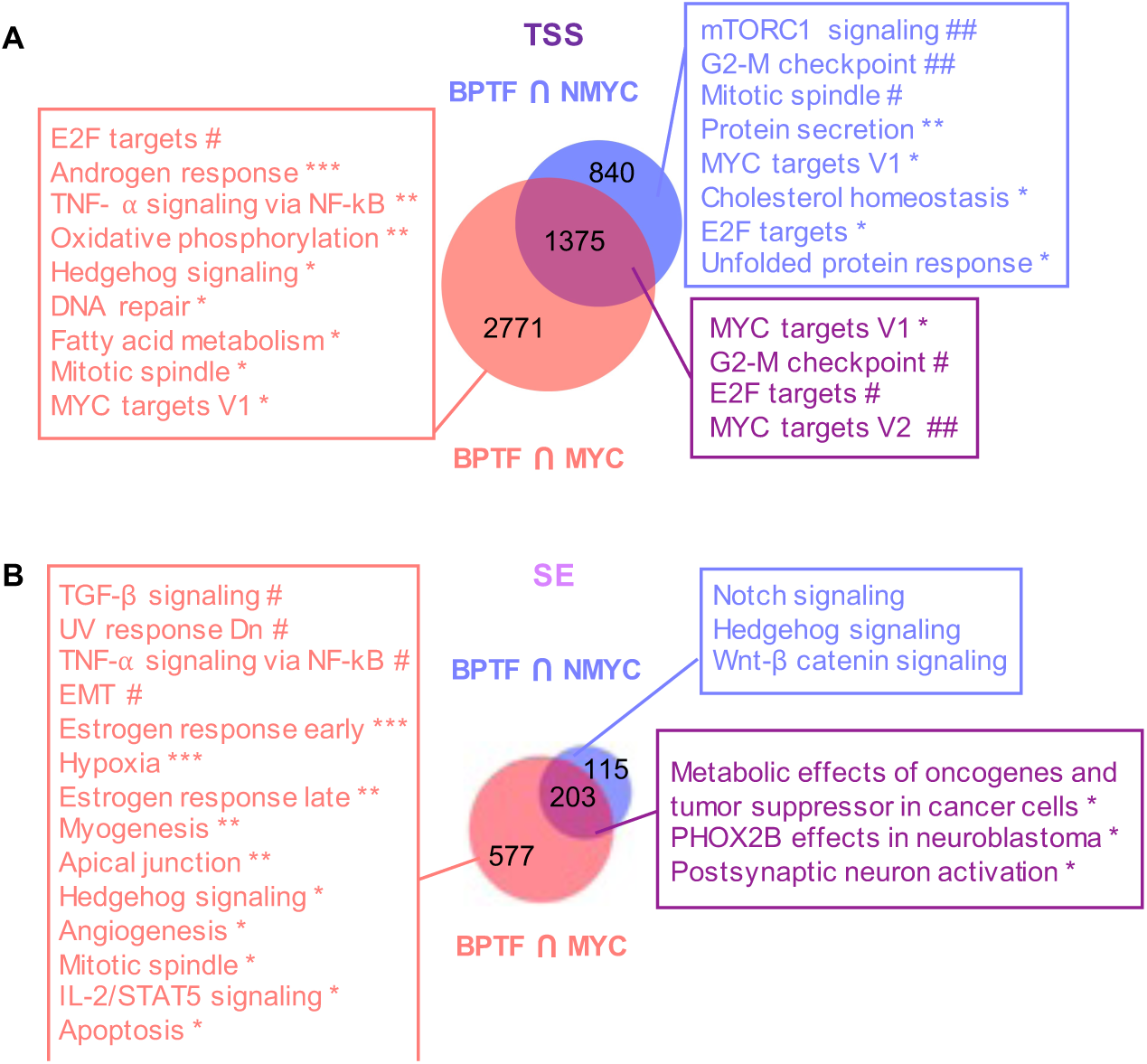
BPTF genomic distribution varies when partnering with MYCN or MYC. (A, B) Comparison of BPTF-MYCN and BPTF-MYC annotated genes in TSS (A) and SE (B). Significantly enriched pathways for each set of genes are shown (blue, BPTF-MYCN only; red, BPTF-MYC specific; purple, both conditions). (*Ad. P* *< 0.05; **<0.005; ***<0.0005; ^#^ <0.00005; ## 0.000005).

### Functional insights on the role of BPTF in neuroblastoma lineage identity

To address the relevance of BPTF to neuroblastoma cell identity, we first assessed the activity of transcriptomic subtype signatures in the SEQC cohort: the ADRN signature activity was significantly higher in the *BPTF*^high^ subgroup while the MES signature was higher in *BPTF*^low^ tumors (**Figure 7A**). Similar enrichments were found in the GMKF and TARGET datasets (not shown). *BPTF* expression levels were also significantly higher in ADRN tumors in the GMKF dataset (adj. *P*=1.5e-06) (**Figure 7B**). We then assessed the overlap of BPTF SE peak-annotated genes with the ADRN or MES signatures ^25^: in KELLY, LAN-1, and KP-N-YN cells the overlap was higher with the ADRN signature; in contrast, in SK-N-AS cells the overlap was higher with the MES signature (**Figure S10A**).

**Figure 7.**
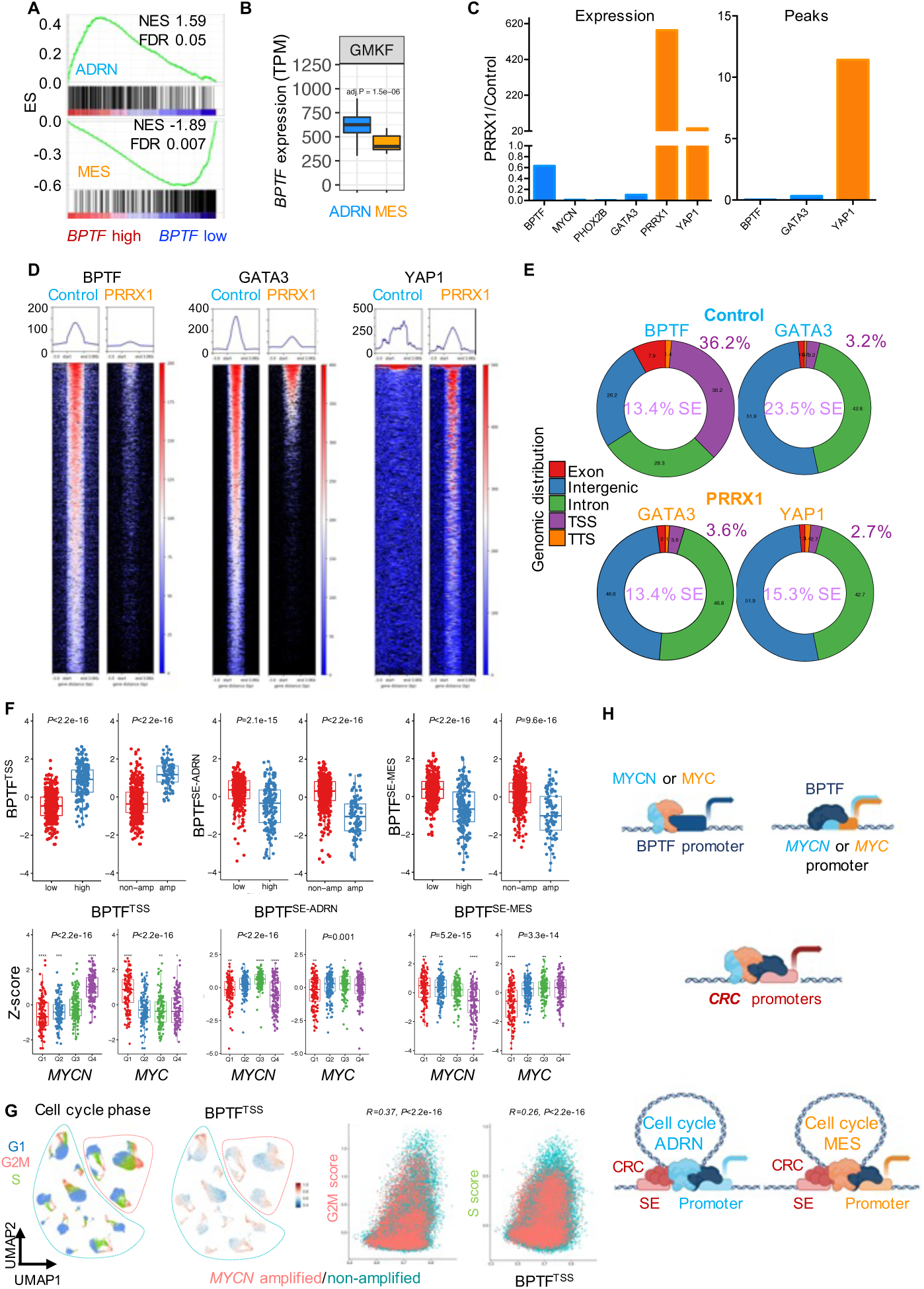
BPTF cooperates with MYC family proteins to link cell cycle and epigenetic states in neuroblastoma cells. (A) The ADRN signature is enriched in *BPTF*^high^ tumors (SEQC dataset, GSEA). (B) Higher *BPTF* expression in ADRN (n=174) than in MES (n=21) tumors from in the GMKF series. (C) Expression of *BPTF*, *MYCN*, *PHOX2B*, *GATA3, PRRX1*, and *YAP1* in KP-N-YN PRRX1-expressing vs control cells (left); ratio of Cut&Run peaks for BPTF, GATA3, and YAP1 in PRRX1-expressing vs control cells (right). (D) Heatmap showing the intensity of the binding of BPTF, GATA3, and YAP1 in Cut&Run experiments performed with control and PRRX1-induced cells. (E) Genomic distribution of BPTF and GATA3 in control cells (left) and GATA3 and YAP1 in PRRX1-expressing cells (right). (F) ssGSEA analysis of BPTF Cut&Run-derived signatures for TSS, SE-ADRN, and SE-MES in the SEQC dataset (z-score adjust). Graphs representing the Z-score value of BPTF Cut&Run-derived signatures according to risk and *MYCN* status (top). Graphs representing the Z-score values of BPTF Cut&Run-derived signatures in relationship to expression of *MYCN* and *MYC*, represented as quartiles. (Kruskal-Wallis test). (G) Cell cycle phase scoring (left) and BPTF^TSS^ binding signature (right) in neuroblastoma cells and correlation between G2/M score and S score with the TSS signature ^32^. (H) Model depicting the role of BPTF in neuroblastoma. MYCN/MYC and BPTF cross-regulate each other (upper panel). BPTF cooperates with MYCN/MYC to regulate the expression of the CRC components (middle panel). BPTF cooperates with MYCN/MYC and with the CRC components to regulate cell cycle genes and the ADRN or MES programs.

To acquire functional insight, we used the inducible PRRX1-mediated ADRN-MES transdifferentiation model in KP-N-YN cells ^22, 25^, using YAP1 as a readout of the MES phenotype. Overexpression of the mesenchymal transcription factor PRRX1 reprograms the SE and mRNA landscapes of ADRN cells toward a mesenchymal state. Upon PRRX1 induction, there was a marked reduction of *BPTF* mRNA levels (ca. 40%) and a dramatic decrease of *MYCN*, *GATA3*, and *PHOX2B* mRNA expression levels (**Figure 7C**). In contrast, *PRRX1* and *YAP1* mRNA levels were strongly up-regulated (580- and 35-fold, respectively) (*P<*0.00001). Cut&Run experiments for BPTF, GATA3, and YAP1 proteins in both conditions showed a marked decrease in the number of BPTF and GATA3 peaks (2% and 30% of control, respectively) in PRRX1-overexpressing cells (**Figure 7C, D**). The very low levels of MYCN in induced MES cells may explain the massive loss of BPTF peaks despite modest changes in expression levels. In contrast, the number of YAP1 peaks was 11.4-fold higher in induced cells (**Figure 7C, D**). Representative peaks are shown in **Figure S10B**. As described above, BPTF binding was more prominent in promoter-TSS regions in control KP-N-YN cells; binding in PRRX1-induced cells could not be robustly assessed due to the low number of peaks called (**Figure 7E**). The genomic distribution of GATA3 was similar in both conditions but the percentage of peaks annotated to NB-SE was higher in control cells (23.5% vs 13.4%, P=0.001). Upon PRRX1 induction, 2.7 % and 15.3% of the YAP1 peaks were annotated to promoter-TSS and SE regions, respectively. Motif enrichment analysis of GATA3 and BPTF peaks showed that the NFY and GATA motifs were enriched in the TSS regions and the GATA motif was significantly enriched in SE-annotated peaks bound by GATA3 both in control and in PRRX1-induced cells (**Figure S7C and S10C**). In contrast, the PHOX2A and AP-1 motifs were differentially enriched in GATA3 peaks in control and induced cells, respectively. Regarding YAP1, the TEAD motif was among the top enriched ones in SE-peaks from PRRX1-induced cells, as was the AP-1 motif (**Figure S10C**). Genes annotated to GATA3 SE peaks in control cells were enriched in neuroblastoma-related pathways (**Figure S10D**), whereas those annotated in PRXX1-induced cells were enriched in EMT and TNF-α signaling via NF-κB), in agreement with a 30% overlap of GATA3-annotated genes in control and PRRX1-induced cells (**Figure S10E**). The significant enrichment of these pathways in genes annotated to YAP1 in induced cells supports a cooperation between GATA3 and YAP1 (**Figure S10F**). BPTF and GATA3 were found to consistently bind to the ADRN CRC genes in KP-N-YN control cells and GATA3 was also found to bind BPTF and its own SE in the induced state. BPTF binding to the ADRN CRC genes was reduced/abolished upon PRRX1 induction (**Figure S10G**).

**Figure S10.**
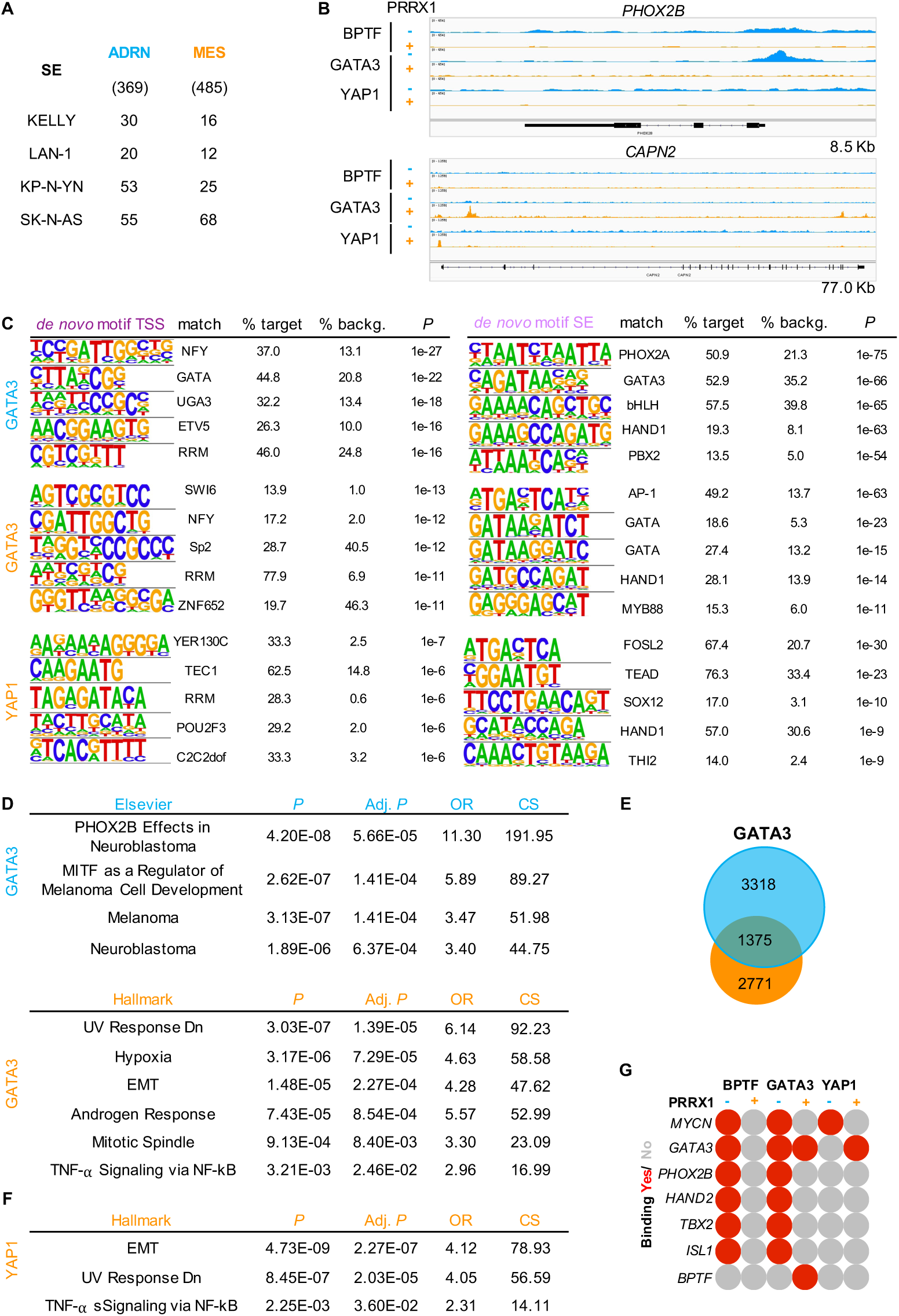
PRRX1-induced ADRN to MES switch results in changes in the genomic distribution of GATA3 and YAP1 (KP-N-YN cells). (A) Overlap of BPTF SE-annotated genes with ADRN or MES signatures. (B) Representative IGV image for BPTF, GATA3 and YAP1 binding to one ADRN gene (*PHOX2B*) and one MES gene (*CAPN2*) in control and PRRX1-overexpressing cells. (C) *De novo* top 5 motifs in the TSS-(left) and SE-(right) annotated peaks (HOMER) for GATA3 (blue, control; orange, induced) and YAP1 (induced). (D) Overlap of pathway analysis for GATA3 SE-annotated genes (blue, control; orange, induced). (E) Overlap of GATA3-annotated genes in control versus PRRX1-induced cells. (F) Significant pathways from the analysis of YAP1-annotated genes in cells overexpressing PRRX1. (G) Binding of BPTF, GATA3, and YAP1 to the genes coding for CRC components in control and induced conditions.

To address the relevance of these findings in the context of patient tumors, we examined the overlap of BPTF-peak gene sets with other tumor-derived signatures. The BPTF^TSS^ signature corresponds to genes annotated to TSS-promoter peaks in KELLY, LAN-1, KP-N-YN, and SK-N-AS cells; the BPTF^SE-ADRN^ signature pertains to genes annotated to SE peaks in at least 2 ADRN cell lines and the BPTF^SE-MES^ refers to BPTF SE peaks in SK-N-AS cells. We compared these gene sets to the SE signatures reported by Gartlgruber et al. in four tumor subgroups (*MYCN*-amplified, high-risk *MYCN*-non-amplified, low-risk *MYCN* non-amplified, and MES) (**Figure S11A**). The overlap of the BPTF^SE-ADRN^ signature was highest with that of SE from *MYCN*-amplified tumors and the overlap of the BPTF^SE-MES^ was highest with the mesenchymal SE signature. In accordance with these findings, the BPTF^SE-ADRN^ and the BPTF^SE-MES^ signatures showed highest overlap with the transcriptomic ADRN and MES signatures reported by Yuan et al. ^37^ (**Figure S11A**), respectively. This was also the case at single cell level (**Figure S11B**). ssGSEA of the BPTF activity score signatures in the SEQC dataset showed that the BPTF^TSS^ signature was highly enriched in *MYCN*-amplified high-risk tumors (**Figure 7F**, P<2.2e-16) expressing higher *BPTF* levels (**Figure 2A**). In contrast, the BPTF^SE-ADRN^ signature was enriched in *MYCN*-non-amplified samples (**Figure 7F**) and the BPTF^SE-MES^ signature was higher in high MYC-expressing tumors (**Figure 7F**). The BPTF^TSS^ signature correlated with G2/M and S phase cell cycle scores in single tumor cell data (R=0.37, p < 2.2e−16; R = 0.26, p < 2.2e−16, respectively) (**Figure 7G**), supporting again the relevance of BPTF in this process. As expected, there was a positive correlation of the BPTF^TSS^ signature with the ADRN-related genes (e.g., GATA3, PHOX2B, HAND2), particularly with *MYCN* (R=0.56, P<2.2e-16) (**Figure S11C**); the BPTF^SE-ADRN^ signature was highly correlated with *HAND2* and *DBH* expression (R=0.38, P<2.2e-16; R=0.68, P<2.2e-16; respectively). The BPTF^SE-MES^ signature activity was also positively correlated with the expression of the MES transcription factors (e.g., PRRX1, NOTCH3), the maximum R value being with *NOTCH3* (R=0.78, P<2.2e-16) (**Figure S11C**).

Collectively, these findings indicate that BPTF plays a key role in the establishment of both the ADRN and MES epigenetic states.

**Figure S11.**
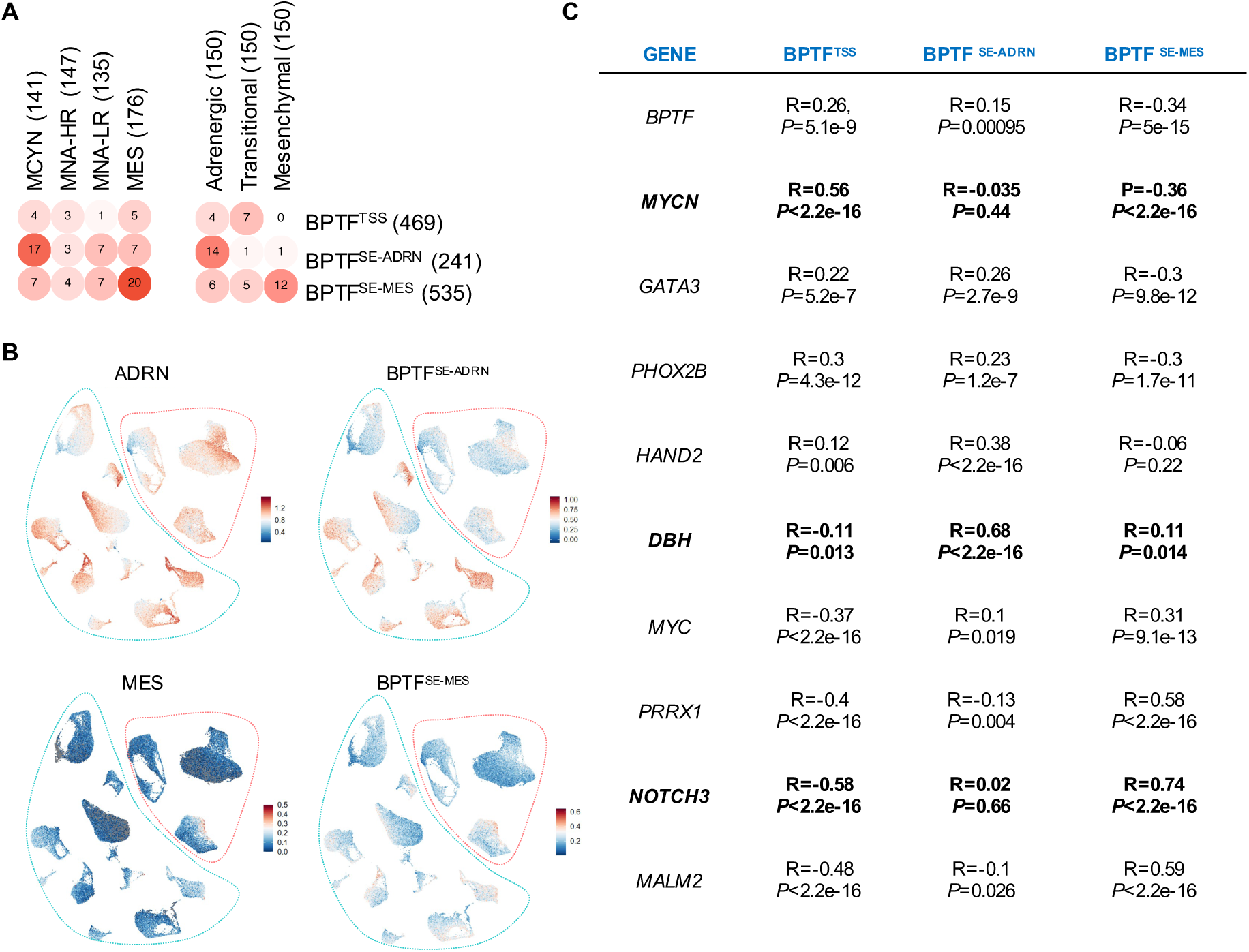
BPTF genome binding signatures from cultured cells are relevant to patient tumors. (A) Heatmap representing the overlap of BPTF Cut&Run-derived signatures with patient tumor-derived SE (left) and transcriptomic (right) signatures ^34, 37^. (B) UMAP graphs showing the ADRN and MES signatures ^25^ and the new BPTF-SE signatures in tumor cells from ^32^. (C) Correlation between expression of *BPTF*, *MYCN/MYC*, and the main ADRN/MES regulators with Z-score values for each BPTF Cut&Run-derived signature (Spearman R).

## DISCUSSION

The requirement of BPTF for the activity of MYC in several cancers, together with the critical role of MYCN in neuroblastoma, led us to explore its role in this tumor. We report that the high expression levels of BPTF in neuroblastoma, among the highest in all cancers, are associated with chromosome 17q copy number gain and with the transcriptional programs driven by MYC family proteins both in *MYCN*-amplified and non-amplified tumors. High *BPTF* expression is associated with high-risk clinical and biologic features. The strong prognostic effect of *MYCN* amplification may underlie the fact that BPTF expression levels are not an independent predictor of outcome. Both *MYCN* and *BPTF* undergo copy number gains in neuroblastoma, possibly contributing to amplify a cooperative regulatory loop playing a key role in the biology of this tumor.

To elucidate the role of BPTF at a mechanistic level, we conducted multilevel genomic analyses using neuroblastoma cell lines as models of the ADRN and MES tumor epigenetic states. Our studies revealed a cooperation of BPTF with both MYCN and MYC. As *MYC* is highly expressed in neuroblastomas lacking *MYCN* amplifications ^38, 39^, our results support a broad role of BPTF across tumor subtypes. Even though *BPTF* transcript levels were high overall, tumors expressing the highest *BPTF* levels displayed highest cell cycle and DNA repair pathway activity. In contrast, tumors expressing lower *BPTF* levels displayed higher activity of inflammatory, NF-κB and EMT related pathways. These findings are reminiscent of our previous findings in murine B cell lymphomas lacking one *Bptf* allele, which displayed higher NF-κB pathway activity than the corresponding tumors from control mice. These observations suggest that overactivation of the NF-κB pathway may contribute to overcome the effects of *BPTF* down-regulation and support cell survival. The findings are also in agreement with other work suggesting that BPTF may participate in EMT ^40, 41, 42^.

To understand how BPTF contributes to the epigenomic landscape of neuroblastoma, we used Cut&Run to acquire information on its genome-wide distribution in cells representative of the main tumor subtypes. This was integrated with genome-wide analysis of the distribution of MYCN/MYC and key components of the ADRN and MES regulatory circuits. As expected, in both ADRN and MES cells, BPTF was found to bind mainly to open chromatin regions with a predominance at promoters, consistent with the notion that it provides specificity to NURF through its interaction with histone modifications (H3K4me3 and H4K16ac) ^43^. Comparison of our Cut&Run data with a recent report on the genomic distribution of BPTF in AML cells, where BPTF preferentially binds to enhancers, active promoters, and insulators, revealed a substantial overlap (approximately 30% of shared regions) ^44^. At promoters, BPTF regulates mainly the expression of cell cycle-related genes in cooperation with MYC family proteins. This cooperation is supported by our previous work demonstrating the presence of BPTF and MYC in the sample complex, now extended to an interaction with MYCN in neuroblastoma cells. Our bioinformatics analyses provide novel evidence that additional transcription factors collaborate with BPTF at promoters, including E2F, NFY, and SP1/2 whose motif binding sites are enriched in BPTF peaks and have been shown to be active in genes regulated during the G2/M transition ^45^. Consistently, a recent report shows cooperation of NFYa and SP2 in activating genes linking metabolism and proliferation in cardiomyocytes

The direct correlation of the expression of BPTF with the ADRN CRC components at the transcript and protein levels observed in our studies prompted addressing whether both types of proteins would be part of the same molecular complex. Using IP-WB and IP-MS we confirmed the presence of BPTF, MYCN, and several CRC components in immunoprecipitates. An interaction of BPTF and MYCN has been previously described in SH-EP cells overexpressing HA-tagged MYCN through lentiviral transduction ^47^; however, all of our experiments were carried out using immunoprecipitated endogenous proteins providing strong biological plausibility to our findings. As expected, several members of the NURF complex, such as BAP18 and RBBP4, were also identified among the BPTF interactors. SMARCA1 and SMARCA5 proteins were found in KELLY but not in SK-N-AS cells. A recent report on an alternative BPTF-containing NURF complex in AML cells, characterized by the presence of BAP18 and SMARCA5 ^44^, suggest that multiple BPTF-containing NURF conformations exist, with potential functional impact.

Two SE-associated TF networks dominate epigenetic control of neuroblastoma phenotypes, shape intratumoral heterogeneity, and are associated with tumor aggressiveness^25^. Cells can transition between the ADRN and MES states, underscoring the need to better understand the mechanisms involved in these neuroblastoma epigenetic states ^48^, with potential impact on therapeutic strategies. Genome-wide BPTF distribution studies showed binding to SEs, including those annotated to genes of the ADRN subtype such as *DBH*, *GATA3*, *PHOX2A,* and *PHOX2B* ^22^, among others, in cells with an ADRN phenotype. In the human single cell data from Thirant et al., *BPTF* was among the genes differentially expressed in one of the ADRN clusters, together with *PHOX2A*, *PHOX2B*, and *ASCL1* ^49^. In ADRN cells, BPTF binds SE annotated to genes related to the ADRN pathway, where it colocalizes with MYCN and the core ADRN CRC components. In MES cells, BPTF binding is enriched at SE of genes involved in EMT, and TGF-β and TNF-α signaling pathways, characteristic of the MES subtype; and it largely colocalizes with MYC. In the absence of *MYCN/MYC* expression, BPTF genomic DNA binding is dramatically reduced. This is in agreement with previous data showing that *MYCN* silencing alters the genomic binding of G9a and WDR5, two MYC cofactors ^50^. These data suggest that BPTF cooperates with both MYCN and MYC at SE to regulate ADRN and MES cell identity programs, respectively, with an additional contribution of the CRC in ADRN cells.

We found a high overlap of the genome-wide BPTF distribution signatures identified in cell lines with ADRN signatures from *MYCN*-amplified tumors and with tumor MES signatures, highlighting the relevance of our findings to patient samples. We also find a strong association between *MYCN* amplification and/or expression with the BPTF^TSS^ and BPTF^SE-ADRN^ signatures. Additionally, we uncover a significant positive correlation between the BPTF^SE-MES^ signature and expression of *MYC* and other MES transcription factors, such as *PRRX1* and *NOTCH3*.

As summarized in the model shown in **Figure 7I**, our work unveils novel roles of BPTF in regulatory circuits that are critical for the activity of MYCN/MYC and the ADRN and MES transcription factors. First, *BPTF* is regulated by MYC family proteins which, in turn, are regulated by BPTF in a feed-forward loop. At another level, BPTF cooperates with MYC family proteins at gene promoters to regulate expression of cell cycle genes and the ADRN CRC components; at yet another level, BPTF cooperates with MYC family proteins and the ADRN and MES transcription factors at SE to promote high-level activity of the corresponding ADRN and MES gene expression programs to confer neuroblastoma cell identity.

The functional genomic analyses and molecular-clinical associations reported here support a critical multilayer role of BPTF in neuroblastoma cell cycle control across neuroblastoma cell types, as well as in the maintenance of epigenetic states and cell identity. Together with previous work showing that reducing BPTF levels genetically a mere two-fold is sufficient to increase mouse survival, our results support BPTF as a relevant therapeutic target in neuroblastoma. The discovery of BPTF degraders is a promising approach with potential application to a wider variety of human tumors.

## METHODS

### Cell culture conditions and treatments

KELLY, LAN-1, and SK-N-AS human neuroblastoma cell lines were kindly provided by Dr. M. Ramirez (Hospital del Niño Jesús, Madrid, Spain). KELLY and LAN-1 cells were cultured in RPMI (Sigma-Aldrich) supplemented with 10% FBS and 1% sodium pyruvate (NaPyr; Thermo Fisher; 1:100). SK-N-AS cells were grown in DMEM (Sigma-Aldrich) supplemented with 10% FBS and 0.1% non-essential amino acids (NEAA; Thermo Fisher; 100x; 1:1.000). KP-N-YN cells expressing an inducible PRRX1 transgene were a kind gift of K. Stegmaier (Dana Farber Cancer Institute, Boston, MA) to the Maris lab. HEK293-FT cells (purchased from the ATCC) were grown in DMEM supplemented with 10% FBS. Cells were grown at 37 °C with 5% CO_2_. Cells were regularly tested and shown to be Mycoplasma-free.

### *BPTF* knockdown

For constitutive knockdown, Mission shRNAs (Sigma-Aldrich) were used for RNA interference. Two BPTF-targeting shRNAs (shBPTF-1, clone TRCN0000016819; shBPTF-2, clone TRCN0000016820) were selected based on optimal activity and compared with a control non-targeting shRNA (NT).

For inducible knockdown, short hairpin (sh) non-targeting (NT), shBPTF#1 and shBPTF#2 targeting oligonucleotides (**Supplementary Table 3**) were annealed and ligated into the pLKO-TET-On vector digested with *Age*I and *Eco*RI. Transformation was performed in *rec*-deficient Stbl3 E. coli. Plasmids were isolated and purified with NucleoSpin Plasmid DNA, RNA & Protein Purification kit (Macherey-Nagel). Clones were screened by *Xho*I digestion and Sanger sequencing.

Lentiviral infections. Lentiviral particles were produced by combining a packaging plasmid (psPAX2) and an envelope-coding plasmid (pCMV-VSVG) together with vectors containing the desired targeting DNA sequences. HEK 293 packaging cells were transfected using JetPrime Transfection reagent, at a ratio of 1: 0.75: 0.25 for transfer, packaging, and envelope plasmids, respectively. Viral particle supernatants were harvested, filtered, and pooled following two rounds of collection at 48 and 72 h. Neuroblastoma cells were seeded at 50% confluence and treated with hexadimethrine bromide (Polybrene) to increase infection efficiency. Media containing lentiviral particles were added to the cells at a 1:1 ratio; two rounds of infection were performed. Infected cells were selected with puromycin, administered at doses previously determined to be optimal for each cell line (1.5-2.5µg/mL). *BPTF* was inducibly knocked-down with doxycycline (1µg/mL). Cells were collected at different time points (24-96 h) post induction to assess the efficiency of *BPTF* knockdown and expression analysis; the 48h time point was selected for RNA-seq experiments.

siRNA knockdown. Control siRNA 1 (ref. D-001810-01-05) and siRNA 2 (ref. D-001810-02-05) and *BPTF*-targetting siRNAs 1 (ref. J-004025-07) and 2 (J-004025-08) were purchased from Dharmacon (GE Biosciences). Cells were subjected to 50 nM concentrations of siRNA in OPTIMEM-I (Gibco) for 6h using lipofectamine 3000 (Invitrogen). Knockdown was confirmed by RT-qPCR at 24-72h post-transfection.

### MYCi975 treatment

To determine cell viability, Celltiter-Glo solution (Promega, ref. G7573) was added to cells grown in 96-well plates. VICTOR3 Multilabel Plate Reader (PerkinElmer) detects the luminescent signal which is proportional to the amount of ATP present in the cells, an indicator of metabolically active cells. KELLY cells were seeded at 10% confluency and treated with vehicle (DMSO) or MYCi975 (ref. S8906, Selleckchem) (6.5μM, IC_50_) 24h after plating. Cells were collected 24-96h later for analysis.

### Cell quantification and viability assay

The Invitrogen Countess Automated Cell Counter 3 (Thermo Fisher) was used to count cells. To determine cell viability, Cell titer-Glo solution (Promega, ref. G7573) was added to cells grown in 96-well plates and measurements were made in a VICTOR3 Multilabel Plate Reader (PerkinElmer).

### Western blotting

For protein extraction, cells were washed with ice-cold PBS and lysed for 30 min in 1% NP40 buffer (50mM Tris-HCl pH 8.0, 150mM NaCl, 1.0% NP-40) supplemented with phosphatase and protease inhibitors, followed by sonication and clearing by centrifugation. Protein extracts were subjected to electrophoresis in 10% polyacrylamide SDS gels using Tris-glycine-SDS running buffer (Alaos) or in NuPAGE 3-8% Tris-acetate precast polyacrylamide gels (Thermo Fisher), with Tris-acetate SDS running buffer (Thermo Fisher). Proteins were then transferred to 0.45 µm nitrocellulose membranes (GE Healthcare), blocked in Tris buffer saline Tween-20 (TBST) 5% skim milk, and incubated overnight with primary antibodies at 4 °C (see Supplementary Table 4). Membranes were then incubated with horseradish peroxidase-conjugated secondary antibodies (Dako; 1:10.000). Reactions were visualized using either Immobilon Crescendo Western HRP Substrate (Sigma-Aldrich) or Immobilon Western Chemiluminescent HRP Substrate (Sigma-Aldrich) in a ChemiDoc MP Imaging instrument (Bio-Rad).

### Protein and monoclonal antibody production

A cDNA encoding the fragment spanning amino acids 701 to 1073 (present in all isoforms) of the human BPTF was amplified by PCR and subsequently cloned into the pDEST-566 vector using Gateway technology (Thermo Fisher Scientific). The resulting 6x-MBP-hBPTF (701-1073) fusion protein was expressed in the Escherichia coli host strain BL21(DE3) (Thermo Fisher Scientific), and the soluble protein was purified on a HisTrap FF column (Cytiva), followed by buffer exchange on a gel filtration column [HiPrep 26/10 Desalting (Cytiva)] using an AKTA protein purification system (Cytiva). The purified fusion protein was concentrated via ultrafiltration and authenticated through mass spectrometry.

Two Balb/c mice were intraperitoneally injected with 100μg of the MBP-hBPTF fusion protein, along with Complete Freund’s adjuvant (Difco), administered four times at 14-day intervals. A final booster of 150μg recombinant MBP-hBPTF protein was injected intraperitoneally, and three days later, splenocytes were fused following the procedure outlined in ^51^. Hybridoma supernatants were screened using ELISA and HEK-293T cells transfected with the pcmvFLAG-hBPTF plasmid. The two mouse monoclonal antibodies developed against BPTF, named TOR102G (IgG2a, k) and TOR249C (IgG1, k), were cloned using the limiting dilution technique.

All animal experiments were conducted in accordance with the approved experimental protocol by the Institutional Committee for Care and Use of Animals from Consejería de Medio Ambiente y Ordenación del Territorio of the Comunidad de Madrid (Madrid, Spain). The experiments were referenced by number 10/423463.9/19, with corresponding project PROEX274/19.

### Ab validation

The specificity of TOR102G and TOR249C for the endogenous BPTF protein was confirmed by IHC using formalin-fixed paraffin-embedded (FFPE) SK-N-AS cell pellets, before and after lentiviral BPTF knockdown). Strong nuclear and cytoplasmic expression of BPTF was observed in wild type SK-N-AS, while reduced staining was observed upon BPTF knockdown in SK-N-AS cells (**Figure S1**). Similar results were obtained by WB (**Figure 1**).

### Protein interaction assays

#### Co-immunoprecipitation assay (Co-IP)

Total cell lysates (2 mg protein) were incubated with primary antibodies overnight at 4 °C. Protein A/G agarose beads (ref. 4RRPAG-1, ABT) pre-blocked with 5% bovine serum albumin were added to the lysates. Following 2 h incubation at 4 °C with rotation, beads were washed three times with NP-40 lysis buffer, and immunoprecipitated proteins were eluted with SDS sample buffer by boiling for 5 min at 95 °C prior to electrophoresis.

#### Immunoprecipitation followed by mass spectrometry analyses

After immunoprecipitation, proteins were eluted from beads in 2%SDS/100 mM Tris-HCl pH8.0. Then, samples were digested using on bead protein aggregation capture (PAC) with MagReSyn® Hydoxyl microparticles (ratio Protein/Beads 1:5) in an automated King Fisher instrument (Thermo). Proteins were digested 16 h ant 37 °C, with 300 µl of trypsin/LysC in 50 mM TEAB pH 8.0 (Trypsin Gold, Promega and LysC, Wako, protein:enzyme ratio 1:100.). Resulting peptides were speed-vac dried and re-dissolved in 20 uL of 0.1% formic acid. LC-MS/MS was done by coupling an UltiMate 3000 *RSLCnano* LC system to an Orbitrap HF-X mass spectrometer (Thermo Fisher Scientific), (KELLY and SK-N-AS) or to an Orbitrap Exploris 480 mass spectrometer (Thermo Fisher Scientific) (MYC). Samples were loaded onto a trap column (Acclaim™ PepMap™ 100 C18 LC Columns 5 µm, 20 mm length) for 3 min at a flow rate of 10 µl/min in 0.1% FA. Then, peptides were transferred to an EASY-Spray PepMap RSLC C18 column (Thermo) (2 µm, 75 µm x 50 cm) operated at 45 °C, and separated using a 60 min effective gradient (buffer A: 0.1% FA; buffer B: 100% ACN, 0.1% FA) at a flow rate of 250 nL/min. The Orbitrap HF-X mass spectrometer was operated in a data-dependent mode, with an automatic switch between MS and MS/MS scans using a top 12 method (intensity threshold ≥ 2e4, dynamic exclusion of 10 sec, and excluding charges unassigned, +1 and ≥ +6). MS spectra were acquired from 350 to 1400 m/z with a resolution of 60,000 FMHW (200 m/z). Ion peptides were isolated using a 2.0 Th window and fragmented using higher-energy collisional dissociation (HCD) with a normalized collision energy NCE of 27. MS/MS spectra were acquired with a fixed first mass of 120 m/z and a resolution of 60,000 FMHW (200 m/z). The ion target values were 3e6 for MS (maximum IT 25 ms) and 1.0e3 for MS/MS (maximum IT, 54 msec). The Orbitrap Exploris 480 mass spectrometer was operated in a data-independent acquisition (DIA) mode using 60,000 precursor resolution and 15,000 fragment resolution. Ion peptides were fragmented using higher-energy collisional dissociation (HCD) with a normalized collision energy of 29. The normalized AGC target percentages were 300% for Full MS (maximum IT of 25 ms) and 1000% for DIA MS/MS (maximum IT of 22 msec). 8 m/z precursor isolation windows with 2 m/z overlap, were used in a staggered-window pattern from 396.43 to 1004.70 m/z. A precursor spectrum was interspersed every 75 MSMS spectra. The scan range of the precursor spectra was 400-1000 mz.

To analyze the data, DDA raw files were processed with MaxQuant (v 2.0.3.0) (KELLY, SKNAS) using the standard settings against a human protein database (UniProtKB/TrEMBL, June 2021, 20,614 sequences) supplemented with contaminants. Carbamidomethylation of Cys residues was set as a fixed modification whereas oxidation of methionines and protein N-term acetylation as variable modifications. Match between runs option was enabled for transference of identifications. Minimal peptide length was set to 7 amino acids and a maximum of two tryptic missed-cleavages were allowed. Results were filtered at 0.01 FDR (peptide and protein level). DIA raw files (MYC) were processed with DIA-NN (ver 1.8.2) using the library-free settings against a human protein database (UniProtKB/Swiss-Prot, 20,614 sequences). Precursor m/z range was set from 390 to 1010 and all other settings were left at their default values. Protein groups with a library q-Value > 0.01 were filtered out. A protein group intensity table was obtained by summing the precursor quantity values from the resulting filtered table.

Afterwards, the protein intensity files were loaded in Prostar (v1.26.0) ^52^ using the intensity values for further statistical analysis. Briefly, proteins with less than 70% valid values were filtered out. Then, a global normalization of log2-transformed intensities across samples was performed using the LOESS function. Differential analysis was done using the empirical Bayes statistics Limma. Proteins with a *P*< 0.05 and a log_2_ ratio >0.5 were defined as enriched. The FDR was estimated to be below 5% by Benjamini-Hochberg correction for multiple testing. Functional analysis was performed using the Perseus software platform ^53^. STRING (https://string-db.org) was used for protein-protein interaction network. The edges indicate both functional and physical protein associations, and line thickness refers to the strength of data. Interaction score=0.9 (highest confidence), k-means clustering for 3 clusters. Edges between clusters marked by dotted line.

### Immunofluorescence staining and PLA

IF analyses were performed on µ-slide 8 wells (ibidi) pre-treated with poly-lysine. Cells were fixed with 4% PFA for 10 min. After washing with PBS, samples were blocked with 2% BSA 0.5% Triton for 1h before incubating overnight with primary antibodies diluted in 1% BSA and 0.25% Triton. Cells were then incubated with A555 anti-mouse and A488 anti-rabbit (Thermo Fisher; 1:200) secondary antibodies for 45 min. After washing with PBS, DAPI was applied for 5 min for nuclear counterstaining before visualization using confocal microscopy (Leica TCS SP5 MP). PLA was performed usin the DuolinkII fluorescence system (Olink Bioscience, Uppsala, Sweden), following manufacturer’s instructions.

### Immunohistochemical analysis of tissue microarrays

All human samples were obtained after informed consent and approval of the corresponding Institutional Ethics Committee.

CHOP series - Tumor samples from 58 neuroblastoma patients were analyzed; 50% of the patients were in the high-risk group. Samples were obtained at diagnosis (55%), post-treatment (38%), or at relapse/metastasis (7%). Samples from 24 PDXs were also analyzed, 67% of which corresponded to *MYCN* amplified tumors.

IBiS series - TMA sections were provided by the Biobank of Hospital Universitario Vírgen del Rocío-Instituto Biosanitario de Investigación de Sevilla (IBiS), Sevilla, Spain). Tumor samples from 49 neuroblastoma patients were used to construct TMAs; 61% of the patients were in the high-risk group. Samples were obtained at diagnosis (49%), post-treatment (24%), or at relapse/metastasis (27%).

Immunohistochemical analyses were performed on TMA sections. Antigen retrieval was performed by boiling for 10 min in citrate buffer pH 6 after deparaffinization and rehydration. Endogenous peroxidase was inactivated with 3% H_2_O_2_ in methanol for 30 min at RT. Sections were incubated for 30 min at RT with 3% BSA 0.1% Triton/PBS, followed by ON incubation at 4 °C with primary antibodies (**Supplementary Table 4**). After washing, the Envision secondary reagent (Dako) was added for 30 min at RT, and sections were washed again three times with PBS. Diaminobenzidine tetrahydrochloride (DAB, ref. SK-4103, Vector Laboratories) was used as a chromogen. Sections were lightly counterstained with hematoxylin, dehydrated, and mounted. Histological images were acquired with a Nikon TE2000 microscope.

Immunostaining of TMAs was evaluated by a pediatric oncology pathologist (J.P.) using digital analysis with QuPath software version 3.2. Following positive cell detection with mean nuclear staining for each antibody, a cell classifier was trained to identify tumor using representative images from all the neuroblastoma tissue microarrays. All TMAs were manually reviewed by the pathologist to ensure accurate classification and to exclude cores with insufficient tumor quantity and presence of necrosis or artifacts. QuPath software was used to generate an H score for the tumor component in each core. The CHOP and IBiS sets of TMAs were analyzed using the same thresholds for positive cell detection.

### Quantitative RT-PCR

RNA was extracted using the ReliaPrep RNA kit (Promega, ref. Z6012) following manufacturer’s instructions. RNA concentration was determined by Nanodrop ND-1000 Spectrophotometer (Thermo Fisher). RNA samples were reverse-transcribed using random hexamers and TaqMan Reverse Transcription reagents (Thermo Fisher, ref. N8080234) with reverse transcriptase in 96 Well Fast Thermal Cycler (Applied Biosystems, ref. 4375305). For qPCR, cDNA samples were then combined with GoTaq qPCR Master Mix SYBR Green (Promega, ref. A6002) in 384 well-plates, analysed in QuantStudio 6 Flex instrument (Applied Biosystems, ref. 4485691). RNA levels were normalised to endogenous *HPRT* using ΔΔCt method. The sequences of the primers used for RT-qPCR are shown in **Supplementary Table 3**.

### RNA-seq library preparation and analysis

RNA from *BPTF* inducible knockdown experiments was isolated as described previously. Average sample RNA Integrity was assayed on an Agilent 2100 Bioanalyzer. Total RNA samples were converted into cDNA sequencing libraries with the “QuantSeq 3‘m RNA-seq Library Prep Kit (FWD) for Illumina” (Lexogen, Cat.No. 015). Briefly, library generation is initiated by reverse transcription with oligodT priming, followed by a random-primed second strand synthesis. The resulting purified cDNA library was applied to an Illumina flow cell for cluster generation and sequenced on an Ilumina NextSeq 550 by following manufacturer’s protocols. Analyses were performed using Nextpresso RNAseq pipeline ^54^. Briefly, sequencing read quality and cross-contamination analysis were performed by FastQC and FastqScreen softwares, respectively. Raw reads were preprocessed using BBDuk software in order to trim polyA and trueseq adapters contaminant sequences and aligned to hg19 human genome assembly using Bowtie and TopHat aligners. Gene counts were generated from alignment maps using htseq-count function. Differential expression analysis was performed using DeSEQ2 R package.

Pathway analyses of differentially expressed genes were performed using molecular signature dataset of GSEA (Gene Set Enrichment Analysis, www.broadinstitute.org/gsea) and Enrichr (https://maayanlab.cloud/Enrichr/). Pearson correlation of data from different samples was calculated from the expression value (expressed as fragments per kilobase of transcript per million mapped reads) of each gene for each sample by using the ‘cor’ command in R (https://www.r-project.org/). Principal component analysis was performed using the ‘prcomp’ command in R, from the correlation value of each sample RNA-seq data from cell lines was downloaded from the Cancer Cell Line Encyclopedia (https://sites.broadinstitute.org/ccle). The human SEQC dataset ^30^ (GSE62564; n= 498 samples) was obtained from the National Centre of Biotechnology Information (NCBI) Gene Expression Omnibus (GEO) (https://www.ncbi.nlm.nih.gov/geo/)); FPKM data from Therapeutically Applicable Research to Generate Effective Treatments (TARGET) and Gabby Miller Kids First (GMKF ) series were downloaded from dbgap (127 high-risk samples) and https://ega-archive.org/studies/phs001436, respectively. The clinical information relevant for this study is summarized in Error! Reference source not found. RNA-seq data were analyzed using GSEA software to compare the subgroups with highest and lowest *BPTF* mRNA decile expression levels, respectively; pathways enriched in each group were identified. Significantly enriched gene sets were those with nominal *P* <0.05 and FDR *q* value <0.25. Promoter Scanning Analysis of up- and down-regulated genes was performed with HOMER. Single-sample GSEA (ssGSEA) was performed using the GenePattern software (https://cloud.genepattern.org/) (v3.9.11_b270). Heat maps representing ssGSEA z-score adjusted enrichment were obtained using Morpheus (https://software.broadinstitute.org/morpheus/).

### chr17q coverage

RNA expression data from TARGET and GMKF cohorts were correlated with coverage values in chr17q region. The coverage values for chr17q region are calculated by taking the ratio of depth of coverage of exons in chr17q region divided by the depth of coverage of all exons in the genome. The correlation is calculated using ggpubr (v0.6.0) package’s *stat_cor()* function and visualized using *ggplot()* function from tidyverse (v2.0.0) R package in R version 4.2.3.

### Methylation beta-value

The beta-value statistic is a number between 0 and 1 where ‘0’ indicates all copies of the CpG locus in the sample are completely unmethylated (hypomethylation) and a value of ‘1’ indicates that every copy of the CpG locus is methylated (hypermethylation) ^31^.

### ChIP-seq and ATAC-seq

ChIP-seq datasets of CRC members (PHOX2B, HAND2, GATA3, MYCN, ISL1, and TBX2) in KELLY cells and MYC in SK-N-AS were obtained from the GEO database (GSE94824, GSE94782, and GSE138295). ChIP-seq datasets of histone marks (H3K4me3, H3K4me1, H3K27ac, and H3K27me3) for both cell lines were obtained from GSE138314. To visualize signal enrichment of ChIP-seq marks, deepTools (v3.5.2) package’s *computeMatrix()* function was used to compute signal distribution across *BPTF* gene, promoter and super-enhancer regions. The matrix with signal values was then provided to *plotProfile()* function with option *-plotType heatmap* to generate heatmaps displaying signal enrichment across lines (profileplyr v1.14.1). KELLY and SK-N-AS ATAC-seq data were downloaded from GEO database (GSE136279).

### Cut&Run genomic binding analysis

Cut&Run experiments were performed as described in ^55^ and ^56^. Briefly, 500,000 cells/sample were harvested, bound to concanavalin A–coated magnetic beads, and permeabilized with 5% digitonin. After overnight incubation, beads were washed and incubated with pAG-MNase (Protein Production Unit, CNIO). Cells were chilled to 0 °C and digestion was initiated by adding Ca^2+^. Reactions were stopped by chelation including spike-in DNA and the DNA fragments released into solution by cleavage were extracted from the supernatant. Library preparation and paired end sequencing for KELLY (5 biological replicates for BPTF; 1 replicate for H3K27me3 and H3K4me3), LAN-1 (4 biological replicates), and SK-N-AS (3 biological replicates) cells were performed at the CNIO Genomics Unit; experiments with the KP-N-YN inducible cells (4 biological replicates) were performed at the Center for Applied Genomics (CHOP).

All replicates of BPTF genome distribution Cut&Run experiments were analyzed using Cut and Run Nextflow pipeline (10.5281/zenodo.5653535). Briefly, reads were alignment to hg19 reference genome and to S. cerevisiae as spike-in genome using Bowtie2 aligner; duplicates were removed using Picard. Peak calling was performed using MACS2 software with mnase-treated chromatin as control sample. Only FDR-significant peaks <0.05 with signalValue > 3 were kept for analysis. Peaks were merged using mergePeaks HOMER function. For down-stream analyses, regions found in at least 3 (KELLY) or 2 (LAN1, SK-N-AS, and KPNYN) replicates were used and annotated using the HOMER annotatePeaks.pl command. Super-enhancer (SE) regions were obtained from merging regions described in cell lines ^25^, with those found in tumors ^34^ using HOMER mergePeaks and were used to filter BPTF peaks using the bedtools intersect command. In the case of GATA3 and YAP1 antibodies the same strategy was followed after performing Motif analysis of each replicate as quality control (presence of GATA and TEAD or AP-1 motifs, respectively). For motif analyses performed in TSS-promoted annotated peaks, hg19 refGene TSS regions were used as background to minimize sequence bias.

### Chromatin-state discovery analysis

ChromHMM analysis considered the combinatorial signal of: BPTF Cut&Run peaks (merge of all replicates); ATAC-seq; MYCN (KELLY), MYC (SK-N-AS), H3K4me1, H3K4me3, H3K27ac, and H3K27me3 ChIP-seq. This was done using *BinarizeBed* command. The resulting output was used to learn a chromatin-state model using a multivariate hidden Markov model (HMM) through the *LearnModel* command. The optimal number of states (n=15) was chosen based on biological annotation of state description according to the original report ^36^.

### Single cell RNA-seq analysis

Single cell data were analysed from sparse count matrices. Briefly, Seurat package was used to load individual sample matrices. Samples were merged in a common object and the metadata from Dong et al ^32^ (GSE137804) was used to filter out cells using the same criteria as in the manuscript and to import authors’ cell annotation. Next, data were normalized using both sctransform and log normalization for the top 2000 variable genes. Both normalized objects were stored as two different slots of the same Seurat object. Linear dimensionality reduction was performed using the first 20 principal components (PC) according to elbow plot visualization. Non-linear dimensional reduction (UMAP) and clustering were performed using these PCs. Cluster stability was visualized in a clustree analysis and resolution was set to 0.2. Markers for the different clusters were obtained with the FindAllMarkers function. Functional analysis was performed using VISION R package and the resultant signature scores were incorporated as metadata to the Seurat object. In the single cell dataset, correlation of BPTF expression with CRC members was visualized in tumor cells. Pearson coefficient was estimated separately for *MYCN* amplified/non-amplified tumor cells with non-zero values for the corresponding gene-pair.

### Graphic representations

Expression heatmaps were generated with MORPHEUS (https://software.broadinstitute.org/morpheus/) and R package. Heatmaps for binding scores associations in genomic regions were created using deepTools software. Briefly, signal matrices were generated using computeMatrix() function from bigwig files restricted to the corresponding genomic regions with the -R flag and associated bed files. The resultant matrices were used to generate the plots using plotHeatmap() command. SRplot platform was also used for data visualization and graphing ^57^. GraphPad Prism was used for graphic representations. Model was Created with BioRender.com.

### Statistical analyses

Spearman and Pearson correlation analyses were performed to compare the expression of two genes of interest. Kaplan-Meier plots were used to assess patient survival. Cox proportional hazards regression models were performed for the association of various clinical covariates and *BPTF* expression groups to assess associations with EFS and OS. Error bars represent the 95% confidence intervals and the indicated P values were from log rank test. For descriptive statistics, P values are computed using a two-sided Wilcoxon test or Kruskal-Wallis tests, considered significant at *P*<0.05. R software version 4.0.2 (R Foundation for Statistical Computing) and GraphPad Prism were used for data analysis.

## Data availability

RNA-seq and Cut&Run data have been deposited in GEO. Accession numbers will be available at the time of publication. The mass spectrometry proteomics data have been deposited to the ProteomeXchange Consortium (http://proteomecentral.proteomexchange.org) via the PRIDE partner repository with the dataset identifier <u>pxd048884</u>. All data related to the manuscript are available upon reasonable request.

## Author contributions

Conceptualization, I.F.; J.M.V, L.D.G., L. M., J.M.M, and F.X.R.; Methodology, I.F., J.M.V, J.M.T, and G.V.; Validation, I.F., J.M.V.; Formal analysis, I.F., J.M.V., M.C., S.S.L., J.M.P., F.G., J.P.; Investigation, I.F., L.S.T., K.T., F.G., and D.M.; Resources, J.M.T., G.R., N.K., D.M., and L.M.; Data curation J.M.V, K.P., and A.F.; Writing, I.F., J.M.V., J.M.M., and F.X.R.; Visualization, I.F., J.M.V, and K.P.; Supervision, F.X.R.; Funding acquisition, L.M., J.M.M., and F.X.R.

## Acknowledgments

This work was supported, in part, by grants SAF2015-70553-R, RTI2018-101071-B-I00, and PID2021-128125OB-I00 from Ministerio de Ciencia, Innovación y Universidades (Madrid, Spain) (co-funded by the ERDF-EU) and RTICC from Instituto de Salud Carlos III (RD12/0036/0034) to F.X.R. I.F. was recipient of a Postdoctoral Fellowship from Juegaterapia-Amigos del CNIO and was partially supported by Asociación NEN, Asociación Pablo Ugarte and Fundación Oncohematología Infantil. CNIO is supported by Ministerio de Ciencia, Innovación y Universidades as a Centro de Excelencia Severo Ochoa SEV-2015-0510. This work was also supported by a National Cancer Institute (NCI) grants R35 CA220500 (J.M.M.) and P01 CA217959 (J.M.M.), and the Giulio D’Angio Endowed Chair (J.M.M.).

We thank Enrique de Alava, Kari Brown, Stephen Chanock, Eduardo Caleiras, Carolina Castilla, Adam Durbin, Elvira Fernández-Vigo, Scott Lowe, Lorena Maestre, Núria Malats, Natalia del Pozo, Manuel Ramírez, Sara Rodríguez, and the Confocal CNIO Unit for valuable contributions; and Luciano di Croce, Juan Méndez, Núria Malats, Laia Richart, Maria Ramal, Mark Kalisz, and other members of the Epithelial Carcinogenesis Group for critical review of the manuscript. J.M.J. thanks Kimberly Stegmaier for providing the inducible KP-N-YN cells.

---

**Supplementary Table 3.**
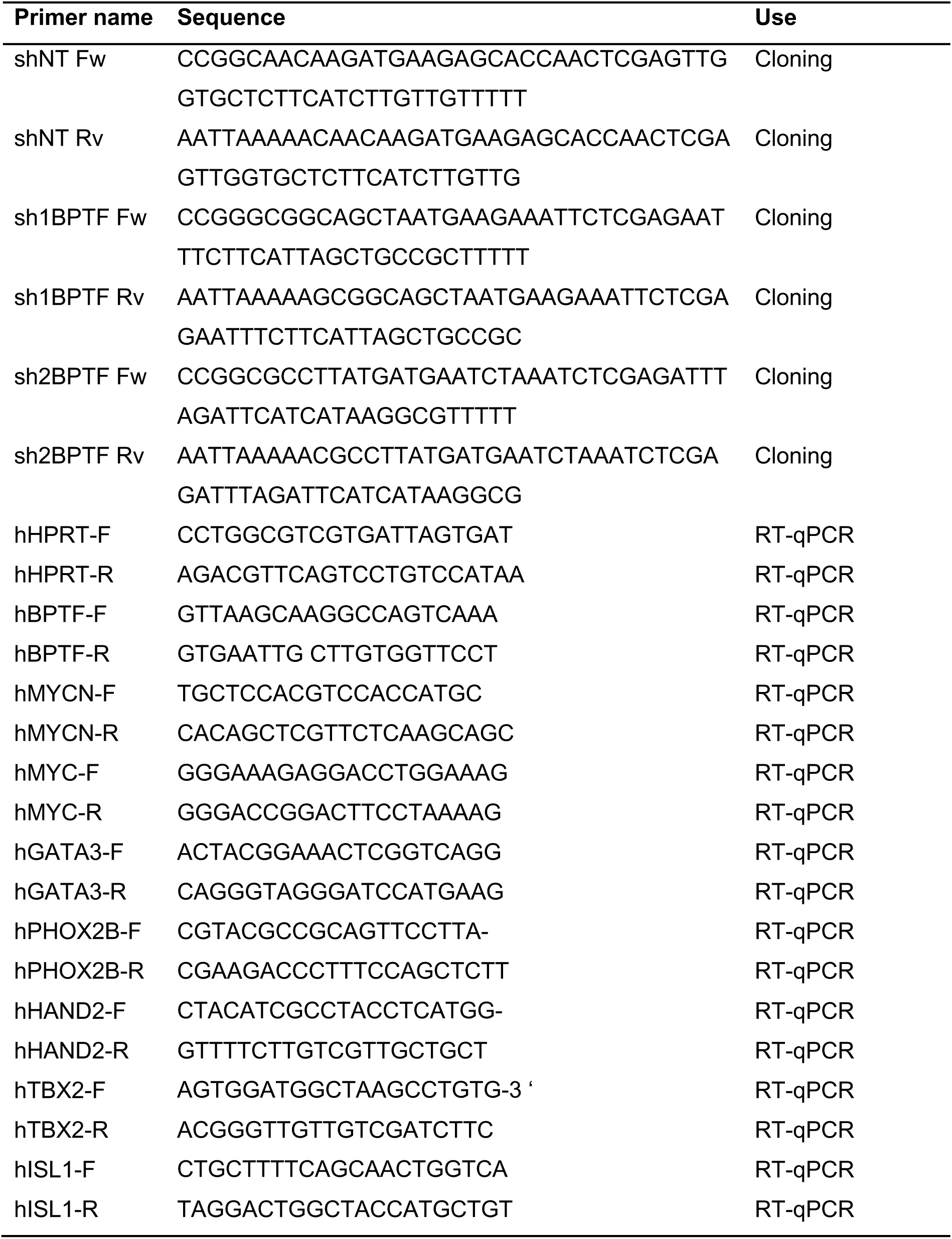
gOligonucleotides used in this study.

**Supplementary Table 4.** List of antibodies used in this study.

| **Antibody** | **Reference** | **Use** |
| --- | --- | --- |
| MYCN | Abcam; ab16898 | WB 1:500 |
| MYC | Abcam, ab32702 | WB 1:10000 |
| BPTF | CNIO Monoclonal antibody Unit;<br>TOR249C/G7 (unpublished) | WB 2 µg/ml |
| BPTF | CNIO Monoclonal antibody Unit;<br>TOR102G (unpublished) | WB 2 µg/ml<br>IP 4 µg |
| BPTF | Sigma-Aldrich-Millipore; ABE24 | WB 1:1000<br>IF 1:100<br>Cut&Run 1:5 |
| BPTF | Sigma-Aldrich-Millipore; ABE1966 | WB 1:1000<br>IHC 1:200 |
| GATA3 | Cell Signaling; GATA-3 (D13C9)<br>XP® | WB 1:1000<br>IHC 1:200<br>IF 1:200<br>Cut&Run 1:50 |
| GATA3 | Biocare medical, CM405A | WB 1:1000 |
| PHOX2B | Santa Cruz, sc-376997 | WB 1:1000<br>IHC 1:100 |
| HAND2 | Santa Cruz, sc-398167 | WB 1:500<br>IF 1:200 |
| ISL1 | DSHB, 39.4D5 | WB 1:1 |
| DAB2IP | Santa Cruz, sc-365921 | WB 1:1000 |
| IQGAP1 | BD Biosciences, BD#610612 | WB 1:5000 |
| YAP1 | Cell Signaling; YAP (D8H1X) XP® | Cut&Run 1:50 |
| H3K23me3 | Cell Signaling; #9377 | Cut&Run<br>1:100 |
| H3K4me3 | Abcam, ab8580 | Cut&Run<br>1:100 |
| Vinculin | Sigma-Aldrich; Clone hVIN-1 | WB 1:10.000 |
| IgG (Mouse) | Santa Cruz, sc-2025 | IP 4 µg |

